# tBid mediated genetic ablation of connective tissue cells reveal their key regulatory function during limb regeneration in axolotls

**DOI:** 10.64898/2025.12.19.695438

**Authors:** Yan Hu, Weimin Feng, Zitian He, Jiayi Zeng, Jingyi Tang, Shulin Li, Tongbo Yu, Sulei Fu, Hui Ma, Binbin Lu, Yanmei Liu, Ji-Feng Fei

**Affiliations:** Guangdong Cardiovascular Institute, Guangdong Provincial People’s Hospital, Guangdong Academy of Medical Sciences, Guangzhou 510080, China; Department of Pathology, Guangdong Provincial People’s Hospital, Guangdong Academy of Medical Sciences, Southern Medical University, Guangzhou 510080, China; School of Basic Medicine, Guangdong Medical University, Dongguan 523808, China; Key Laboratory of Brain, Cognition and Education Science, Ministry of Education, China; Institute for Brain Research and Rehabilitation, and Guangdong Key Laboratory of Mental Health and Cognitive Science, South China Normal University, Guangzhou 510631, China; School of Basic Medical Sciences, Southern Medical University, Guangzhou 510515, China; Guangdong Engineering Research Center of Precision Detection and Modulation of Human Microbiome, School of Life Sciences, South China Normal University, Guangzhou 510631, China; The Innovation Centre of Ministry of Education for Development and Diseases, School of Medicine, South China University of Technology, Guangzhou 510006, China

**Author notes:** These authors contributed equally to this work (Yan Hu, Weimin Feng, Zitian He, Jiayi Zeng). Corresponding authors. *E-mail addresses* (Y. Liu), (J-F. Fei).

**Keywords:** axolotl, cell ablation, connective tissue, limb regeneration, Prrx1

## Abstract

Limb regeneration in adulthood is a fascinating phenomenon unique to salamanders, requiring precise coordination and interaction of various cell types. Connective tissue (CT) constitutes a major component of the limb and serves as the primary reservoir of positional information during regeneration. However, the direct evidence and the extent of CT in positional contribution remain limited. Here we develop an inducible Cre-LoxP-mediated *tBid* cell ablation system in axolotls and demonstrate efficient elimination of targeted muscle tissue and muscle stem cells through transient electroporation or genetic approach. Ablation of CT cells at early stages of regeneration results in delayed regeneration and loss of proximal limb segments (e.g., upper and lower limb), with minimal impact on hand differentiation. CT ablation during development yields similar defects as observed during regeneration, indicating an essential role for CT cells in determining the positional identity of developing and regenerating limb structures. Further scRNA-seq reveal a progressive proximal-to-distal transition among CT cells and identify distinct CT subtypes that contribute to proximal and distal segments. CT ablation results in reduction of the differentiated CT1—a subpopulation that maintains proximal identity throughout regeneration, and a delay of distal transformation in the proliferative CT1—a major CT subtypes showing progenitor property. These cellular shifts likely lead to the observed morphological deficits in regenerating limbs after CT ablation. Moreover, differentiated CT1 cells enhanced their interactions with surrounding cell types after CT ablation, to modulate adaptive proliferation in other populations, e.g., muscle stem cells. Our work establishes a *tBid*-based ablation strategy for functional studies of CT and other cell types in axolotls, and provides direct evidence demonstrating pivotal roles of CT cells in guiding positional memory and segmental patterning during organ regeneration.

## Introduction

Salamanders are the only tetrapod capable of regenerating fully functional limbs as adults^[1–7]^. They can also faithfully regenerate a variety of other complex structures, including parts of the brain, spinal cord, and certain internal organs^[8–13]^. The axolotl is the most widely used salamander species in laboratory and serves as a valuable model in regenerative biology. Limb regeneration involves the coordinated participation of multiple cell types, which either directly contribute progeny to the new tissue or provide essential instructional cues to support the process. While several cell types, such as neurons, macrophages, muscle stem cells, and connective tissue (CT) cells, have been shown to play indispensable roles, others, like muscle tissue, are largely dispensable for regeneration^[14–25]^. However, key questions such as the precise function of many cell types, the extent of their individual contributions, and the molecular interactions that coordinate their behavior during limb regeneration, largely remain unresolved.

CT cells are the most abundant tissue type in the vertebrate limb, exhibiting remarkable morphological and functional heterogeneity ^[24, 26–29]^. In salamanders, CT cells play an integral role in both limb development and regeneration^[24, 26, 28–31]^. Following limb amputation in axolotls, lineage tracing studies have established that CT cells are the primary contributors to the blastema, a proliferative progenitor cell mass essential for regeneration^[24, 28, 29, 32]^. These CT-derived blastema cells reactivate a transcriptional program reminiscent of early limb development, re-expressing key morphogens (e.g., Shh, FGFs, BMPs) and transcription factors (e.g., Hox genes, Tbx, Lmx-1, Hand2)^[4, 14, 33–42]^{Gerber, 2018 #1434}. Ultimately, they differentiate into diverse lineages, including fibroblasts, chondrocytes, osteoblasts, tendons, and ligaments, to reconstitute the regenerated limb.

Beyond providing structural integrity, CT cell function as a critical signaling center that orchestrates blastema formation and patterning. They secrete and respond to essential growth factors and morphogens (e.g., FGFs, BMPs, Wnts) and actively remodel the extracellular matrix to create a permissive microenvironment for regeneration^[1, 29, 39, 43–47]^. Furthermore, CT cells are suggested to be a primary reservoir of positional information, which ensures the accurate re-establishment of limb segments along the proximal-distal axis^[14, 30, 35, 37, 48–50]^. Several molecular cues, including retinoic acid, Shh, FGF, Prod1, Tig1 and transcription factors such as Meis, Hox, Tbx, and Shox2, have been implicated in specifying anterior-posterior, dorsal-ventral, and proximal-distal positional identities^[14, 30, 34, 35, 37, 38, 49, 51–57]^. Recent chromatin profiling studies in axolotl limb CT cells have identified segment-specific levels of histone H3K27me3, particularly at limb homeoprotein gene loci like *Hoxa13*, as a major positional mark encoding segment identity^[58, 59]^. This supports the model wherein positional information is embedded within the CT. Observations that certain CT subpopulations exhibit behaviors consistent with “distal transformation” during regeneration further substantiate their role in positional guidance^[14, 30, 53]^. However, direct evidence elucidating the precise extent of CT’s contribution to positional identity, and the specific mechanisms by which CT cells regulate the activities of other cell types during limb regeneration, remains an area of active investigation.

Accurate elimination of targeted cell types provides a powerful approach for dissecting the roles of specific cell populations and their interactions with other cells during physiological and pathological processes. The removal of specific cell populations, such as CT cells, in a spatially and temporally controlled manner, enables precise determination of their functions during regeneration. To this end, several genetically encoded cell ablation systems have been developed and applied across diverse organisms. These include nitroreductase (NTR)-based methods^[60–64]^, diphtheria toxin (DT) and diphtheria toxin receptor (DTR) systems^[65, 66]^, Kid/Kis^[67, 68]^, thymidine kinase (TK)^[69, 70]^, and *tBid*-based strategies^[71–73]^. The tBid protein is a cleaved form of the “BH3-only” pro-apoptotic protein Bid. Upon induction, tBid translocates from the cytoplasm to the mitochondria, where it binds and activates Bax/Bak, thereby triggering mitochondrial outer membrane permeabilization and apoptosis^[74, 75]^. An inducible *tBid* expression system has been successfully employed to achieve genetic ablation of specific cell types, such as pancreatic β-cells in zebrafish and lens cells in Xenopus, as well as in cultured human cells^[71–73]^. As a component of the endogenous apoptosis pathway, *tBid*-mediated ablation is expected to elicit minimal inflammatory response and off-target toxicity, thereby reducing undesired damage to surrounding tissues.

In this study, we established a *Cre-LoxP* based, *tBid*-mediated conditional cell ablation system in axolotls and investigated the role of CTs during limb development and regeneration. Early CT ablation led to delayed regeneration and loss of proximal limb structures, while hand formation was less affected. Similar defects emerged from early developmental CT ablation, underscoring the consistent importance of CT in positional patterning across life stages. This defect correlated with early *Hoxa* gene expression showing dramatic reduction of the molecular defined proximal but not distal domain. scRNA-seq identified proximal-to-distal transition and dynamics during limb regeneration. And further revealed that CT ablation resulted in reduced number, and delayed distal transformation of certain CT subtypes, which may account for the limb regeneration defects. Our findings not only present a versatile genetic tool for precise cell ablation in the axolotls but also define essential roles for CT in guiding positional memory and segmental regeneration.

## Results

### Establishment of *tBid*-mediated cell ablation in axolotls

To assess whether *tBid* induces cell ablation in axolotls, we first cloned the *pCAGGs:Cherry-T2A-tBid* plasmid in which a constitutive *CAGGs* promoter drive the expression of both CHERRY and tBid protein. We then introduced the mixture of the *pCAGGs:Cherry-T2A-tBid* and *pCAGGs:GFP* plasmids, or the mixture of *pCAGGs:Cherry* and *pCAGGs:GFP* plasmids as control into the tail muscle tissue of the axolotls by electroporation (Figure S1A,B, Supporting Information). It is expected that *tBid* expression would lead to the loss of electroporated cells in the targeted tissues. We observed that at five days post-electroporation, CHERRY fluorescence signal was rarely detected, along with a low level GFP fluorescence in tail after *pCAGGs:Cherry-T2A-tBid* and *pCAGGs:GFP* plasmids electroporation, when compared to controls (Figure S1C, Supporting Information). Immunofluorescence on tissue sections using antibodies against GFP, CHERRY and cleaved Caspase3 (an early apoptotic marker), revealed a rare and sparse distribution of GFP and CHERRY fluorescence signals, co-localized with Caspase3 expression in *pCAGGs:Cherry-T2A-tBid* and *pCAGGs:GFP* plasmids electroporated tails. Conversely, GFP and CHERRY double-positive cells were clearly detected on sections from the controls that lack Caspase3 signals (Figure S1D, Supporting Information). These preliminary results suggest that transient expression of the *tBid* in axolotl leads to ablation of the targeted cells, via apoptosis pathway, in line with previous observations^[71–73]^.

We next evaluated the in vivo performance of *tBid* in transgenic axolotls. To this end, we introduced the plasmid *Tol2-CAGGs:loxp-EGFP-loxp-tBid* together with *Tol2* transposase mRNA into single cell stage axolotl eggs from breeding of *d/d* or *Pax7^Cre-ERT^* ^[23]^, to generate *Caggs^EGFP/tBid^* transgenic line or *Pax7^Cre-ERT^*/*Caggs^EGFP/tBid^* double transgenic (*Pax7-tBid dTG*) line (Figure 1A). The derived F0 *Pax7-tBid dTG* and *Caggs^EGFP/tBid^* axolotls were raised for germline transmission, and the *Pax7-tBid dTG* double transgenic individuals were developed for experiments. In *Pax7-tBid dTG* animals, *tBid* expression is specifically induced in PAX7+ cells upon tamoxifen-induced recombination. Considering the limited number of *Pax7-tBid dTG* F0 axolotls, we carried out blastema transplantation from *Pax7-tBid dTG* donors to *d/d* hosts to generate chimeric axolotls to expand the number of double transgenic limbs (Figure 1B). It is observed that EGFP is ubiquitously expressed in limb transplants before tamoxifen (4-OHT) treatment, indicating their donor origin from *Pax7-tBid dTG* chimeras (Figure 1C). Immunofluorescence on cross sections showed a significant reduction of PAX7+ MuSCs (muscle stem cells) in the transplanted *Pax7-tBid dTG* limbs compared to that from the *d/d* control limbs at 8 days post-treatment (dpt) (Figure 1D). Quantification revealed that the number of PAX7+ cells decreased to 53 per cross section in transplanted limbs, compared to 302 cells in *d/d* limbs at 8 dpt (Figure 1E). Additionally, we tested ablation of PAX7+ cells during forelimb development in germline transmitted F1 *Pax7-tBid dTG* line. Immunofluorescence on longitudinal sections showed that PAX7+ cells were nearly absent in limb buds from *Pax7-tBid dTG* axolotls at 5 days post 4-OHT treatment, compared to controls treated with DMSO (Figure 1F). Collectively, these results demonstrate that the conditional *Cre-LoxP-based tBid* transgenic line generated in this study enables tissue-specific ablation of targeted cells, when combined with an inducible Cre line^[23]^.

**Figure 1.**
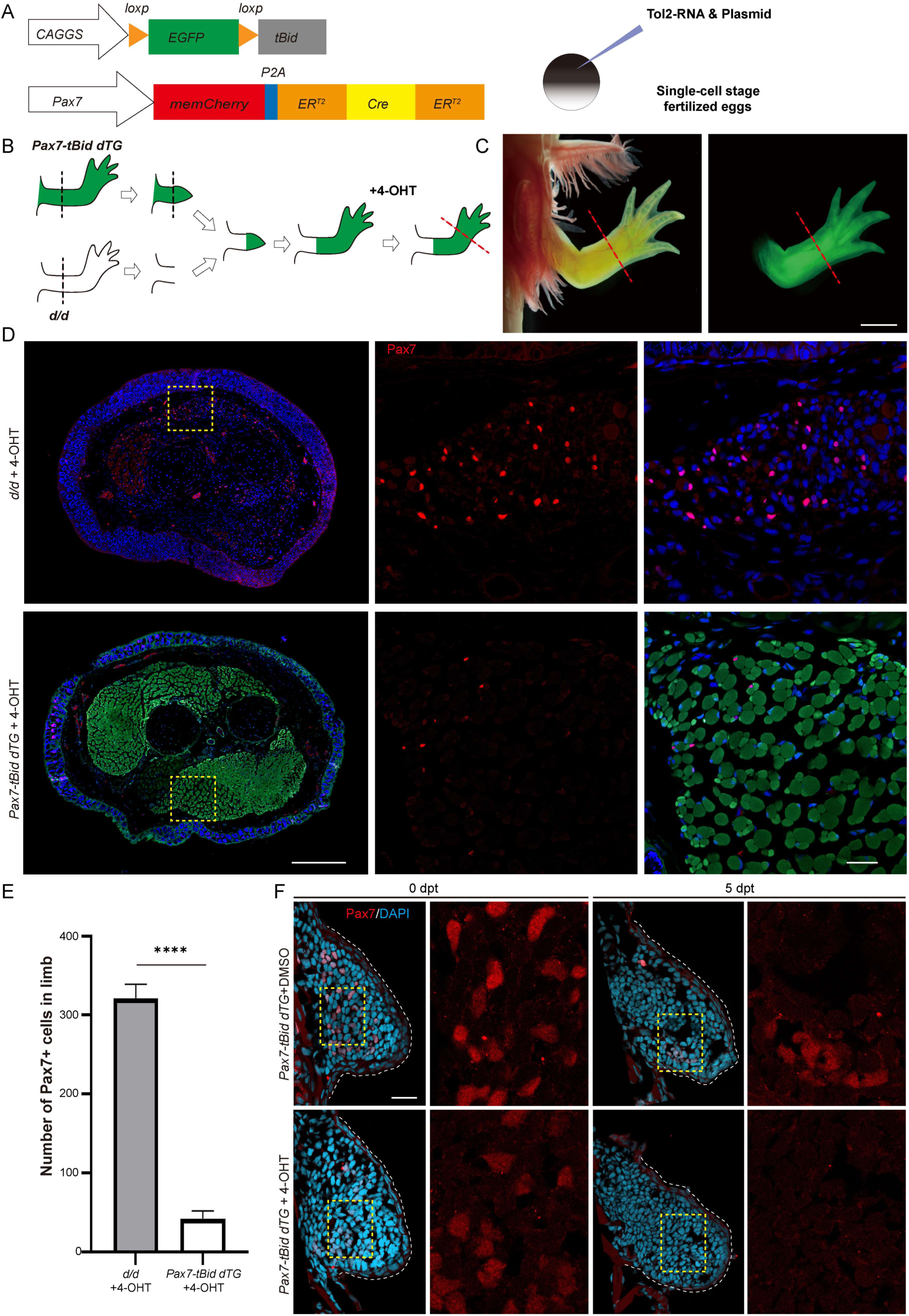
Induced *tBid* expression effectively ablates targeted PAX7+ cells. A) Schematic of the generation of *Pax7-tBid dTG* axolotls. B) Scheme for the muscle stem cell ablation via blastema transplantation. The green limb represents the *Pax7-tBid dTG* donor, the white limb indicates the *d/d* host. C) Live images of bright field (BF) and EGFP fluorescence of the *Pax7-tBid dTG* limbs after blastema transplantation and before tamoxifen treatment. EGFP fluorescence shows the transplanted blastema from a *Pax7-tBid dTG* donor to a *d/d* host at 1 month post-transplantation. D) Immunofluorescence staining for PAX7 (red) and endogenous EGFP fluorescence (green) with DAPI (blue) on limb cross-sections from d/d (upper panels) and *Pax7-tBid dTG* transplanted blastemas (lower panels) at 8 days post-treatment (dpt). PAX7 signal is virtually absent in the transplanted *Pax7-tBid dTG* blastema. E) Quantification of PAX7+ cells in limb tissues at 8dpt (n=3 each). F) Immunofluorescence staining for PAX7 (red) and DAPI (blue) on longitudinal-section of *Pax7-tBid dTG* limb buds at 0 and 5dpt following administration of tamoxifen (4-OHT) or DMSO. PAX7 signal is absent at 5dpt in *Pax7-tBid dTG* axolotls. Black and red dashed lines show the planes of amputation before blastema transplantation, and after tamoxifen conversion, respectively. Areas in the boxes are presented at a higher magnification, showing either single-channel or merged images. White dashed lines outline the shape of limb bud. Data were analyzed by unpaired two-tailed Student’s t-test and represented as mean ± SEM, \*\*\*\**p*<0.0001. Scale bars: 2 mm in (C); 500 μm in (D) left; 50 μm in (D) right; 100 μm in (F).

### Ablation of CT cells during axolotl limb development impairs limb patterning

Previous studies have suggested that CT cells are the potential carrier of the positional and patterning information^[14, 30, 31, 35, 37, 48–50]^. To provide the direct evidence and determine the extent of CT participation during axolotl limb development, we studied the process of limb development after CT cell ablation. Toward this end, we bred *Prrx1^Cre-ERT^* axolotls, in which expression of an inducible Cre is driven by the mouse *Prrx1* promoter^[24]^, with *Caggs^EGFP/tBid^* line to establish *Prrx1^Cre-ERT^*/*Caggs^EGFP/tBid^* double transgenic (*Prrx1-tBid dTG)* line (Figure 2A). Living image revealed that TFP expression from *Prrx1* transgenic allele was confined to the limb bud in the *Prrx1^Cre-ERT^*, while EGFP expression from *tBid* locus throughout the body in *Prrx1-tBid dTG* (Figure 2B) axolotl larvae.

**Figure 2.**
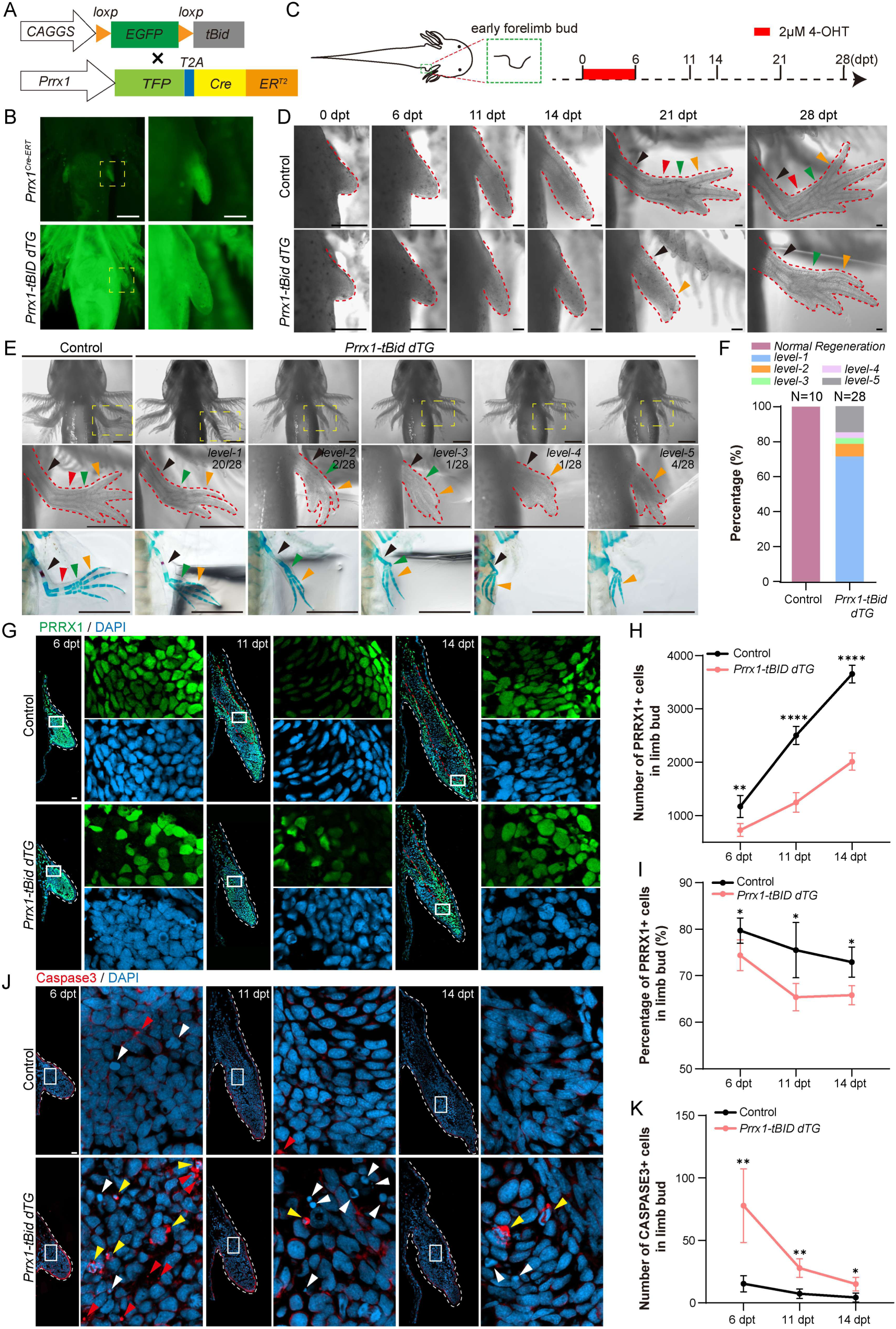
Early ablation of CT cells impairs the development of proximal limb segments, whereas distal hand growth is less affected. A) Schematic representation of breeding between *Prrx1^Cre-ERT^* and *Caggs^EGFP/tBid^* to generate *Prrx1-tBid dTG* axolotls. B) Live images of TFP fluorescence in *Prrx1^Cre-ERT^* (upper panels) and EGFP fluorescence in *Prrx1-tBid dTG* (lower panels) axolotls. TFP expression is restricted to the limb bud in *Prrx1^Cre-ERT^*, whereas *Prrx1-tBid dTG* axolotls exhibit ubiquitous EGFP expression. The right panels show higher-magnification views of the regions outlined by yellow dashed boxes. C) Timeline of 4-OHT administration during early forelimb development. D) Representative time-series images of forelimb development after 4-OHT treatment in control (upper panel) and *Prrx1-tBid dTG* (lower panel) axolotls. Limb development in *Prrx1-tBid dTG* animals was delayed and exhibited defects in the zeugopod segment. E) Live images and corresponding Alcian blue/Alizarin red skeleton staining of forelimb following early 4-OHT treatment. Five distinct phenotypes were observed following CT cell ablation. The middle panels show higher-magnification views of the areas within the yellow dashed boxes. F) Phenotypic analysis with alcian blue/alizarin red skeleton staining of forelimbs following early 4-OHT treatment. G) Immunofluorescence for PRRX1 (green) combined with DAPI (blue) on limb bud longitudinal-sections from control and *Prrx1-tBid dTG* axolotls at 6, 11, and 14 dpt. The *Prrx1-tBid dTG* axolotls show significant reduction in PRRX1+ cells compared to controls. H) and I) Quantification of the absolute number (H) and percentage (I) of PRRX1+ cells in limb bud at 6, 11 and 14 dpt in control (black line, n = 4 each) and *Prrx1-tBid dTG* (red line, n = 4 each) axolotls. J) Immunofluorescence for cleaved Caspase-3 (red) and DAPI (blue) on limb bud longitudinal sections from control and *Prrx1-tBid dTG* axolotls at 6, 11, and 14 dpt. The *Prrx1-tBid dTG* axolotls exhibited a significant increase in cleaved Caspase-3 signal compared to controls. K) Quantification of the absolute number of cleaved Caspase-3+ cells in limb buds at 6, 11 and 14 dpt in control (black line, n = 6 each) and *Prrx1-tBid dTG* (red line, n = 4 each) axolotls. White boxes outline the higher magnification area of limb bud, showing either single-channel or merged images in (G) and (J). Red dashed lines outline the shape of limb in (D) and (E), and shape of skeleton within limb in (G). White dashed lines outline the shape of limb in (G) and (J). Black arrowheads indicate the stylopod; red arrowheads indicate the zeugopod; green arrowheads indicate the autopod; yellow arrowheads indicate digits in (D) and (E). Yellow arrowheads indicate the cleaved Caspase-3-positive cells with nucleus; white arrowheads indicate the ruptured nucleus without the cleaved Caspase-3 signals; red arrowheads indicate the Caspase-3 signals without nucleus in (J). For quantification, data were analyzed by unpaired two-tailed Student’s t-test and represented as mean ± SEM, \**p* <0.05; \*\**p* <0.01; \*\*\*\**p* <0.0001. Scale bars: 500 μm in (B); 1 mm in (E); 100 μm in (D), (G) and (J).

To evaluate the consequences of CT cell ablation on limb development, we first administered 4-OHT to both *Prrx1-tBid dTG* axolotls and the control *Caggs^EGFP/tBid^* at stages 36-38, early forelimb bud stages (Figure 2C and Figure S2A, Supporting Information). Upon 4-OHT administration, excision of the floxed *EGFP* cassette resulted in *tBid* expression driven by the *CAGGs* promoter. At 14 dpt, limb patterning was delayed in *Prrx1-tBid dTG* axolotls, compared to that from the controls showing formed digits. Such delayed distalization phenotype (digit formation) became extremely obvious at 21 dpt. By 28 dpt, all 4-OHT-treated *Prrx1-tBid dTG* animals exhibited forelimb malformations, characterized by varying degrees of developmental delay and consistent loss of the zeugopod (Figure 2D). Phenotypic categorization of the CT-ablated limbs revealed five distinct morphological outcomes (Figure 2E,F): (1) the majority of forelimbs (72%, 20/28) retained a complete stylopod and autopod, including digits, but lacked the zeugopod; (2) 7% (2/28) of forelimbs displayed a complete stylopod and autopod, with loss of the zeugopod and partial digit reduction; (3) 3.5% (1/28) exhibited loss of the zeugopod along with reductions in the stylopod, autopod, and digits; (4) 3.5% (1/28) showed loss of both the zeugopod and autopod, accompanied by a reduced stylopod and digit number; (5) and 14% (4/28) consisted solely of digits, with complete absence of the stylopod, zeugopod, and autopod (Figure 2E,F). Notably, while limb length was significantly reduced in *Prrx1-tBid dTG* axolotls compared to *Prrx1^Cre-ERT^* controls in response to 4-OHT treatment, their overall body size was unaffected (Figure S2B,C, Supporting Information).

We next examined the efficiency of CT cell ablation via immunofluorescence staining of limb bud longitudinal sections at 6, 11, and 14 dpt using an antibody against PRRX1. Compared with controls, *Prrx1-tBid dTG* limb buds displayed reduced size, developmental delay, and a pronounced reduction in PRRX1+ cells (Figure 2G). Corresponding to these morphological abnormalities, quantitative analysis confirmed a significant decrease in the absolute number of PRRX1+ cells at all time points examined (6, 11, and 14 dpt) in *Prrx1-tBid dTG* animals (Figure 2H). Additionally, the proportion of PRRX1+ cells within the limb bud was significantly lower in *Prrx1-tBid dTG* axolotls relative to controls (Figure 2I). Immunofluorescence detection of cleaved Caspase3 further revealed elevated cell death in *Prrx1-tBid dTG* limb buds (Figure 2J). Quantification confirmed a significantly higher number of apoptotic cells in *Prrx1-tBid dTG* limbs at 6, 11, and 14 dpt, with the peak observed at 6 dpt (Figure 2K). Together, these data show efficient ablation of PRRX1+ CT cells via apoptosis upon *tBid* induction and indicate that CT cells are essential for proper patterning and segmentation during early axolotl forelimb development.

To investigate the role of CT during later forelimb developmental stage, we administered 4-OHT at stages 51-52, coinciding with the initial formation of the fourth digit in forelimbs (Figure S3A, Supporting Information). In *Prrx1-tBid dTG* animals, later 4-OHT treated forelimbs exhibited partial loss of the autopod and digits, along with size reduction of the zeugopod compared to controls (Figure S3B, Supporting Information). These results suggest that CT cells are required not only for early limb segmentation but also for the attainment of full morphological complexity at later stages. Given that hindlimbs were nascent limb buds when treated with 4-OHT at stages 51-52, we also assessed the consequences of CT cell ablation on hindlimb development. In *Prrx1-tBid dTG* axolotls, this ablation severely impaired hindlimb development. The resulting limbs exhibited incomplete stylopod formation, a complete absence of the zeugopod and autopod, and malformed and reduced digits (Figure S4A,B, Supporting Information). These defects, which persisted into adulthood (Figure S4B, Supporting Information), are consistent with our earlier observations from CT ablation during early forelimb development (Figure 2D,E). Taken together, our findings demonstrate that CT cells are indispensable for the proper segmentation and patterning of both forelimb and hindlimb development in axolotls, suggesting a conserved regulatory mechanism.

### Ablation of CT cells impairs limb patterning during axolotl limb regeneration

Given that CT constitutes a major cellular component of the blastema and has been implicated as a key contributor to limb regeneration and a potential regulator of positional identity^[24, 26, 28–30, 32, 37, 50]^, we investigated the role of CT cells for forelimb regeneration at stage 51-52 axolotls. To this end, 4-OHT treatment was initiated ten days before amputation and continued until 6 days post-amputation (dpa), for a total of nine administrations (Figure 3A). Monitoring 4-OHT-treated axolotls from 2 to 57 dpa revealed that the limbs of *Prrx1-tBid dTG* animals were significantly smaller than that of controls from 15 dpa onward (Figure S5A,B, Supporting Information). Regenerating forelimbs of *Prrx1-tBid dTG* animals exhibited severe defects compared to controls, characterized by the absence of both the zeugopod and autopod, and the formation of only two to three digits (Figure 3B). Phenotypic analysis of CT cell-ablated regenerating forelimbs revealed two distinct outcome categories: (1) The majority (53%) of regenerates exhibited severe truncation, characterized by the complete absence of the zeugopod and autopod. In these cases, only two or three digits were present, attached directly to a truncated stylopod remnant; (2) The remaining regenerates (47%) were characterized by the absence of the zeugopod, along with incomplete regeneration of the autopod and digits, which were attached to a remnant stylopod (Figure 3C,D).

**Figure 3.**
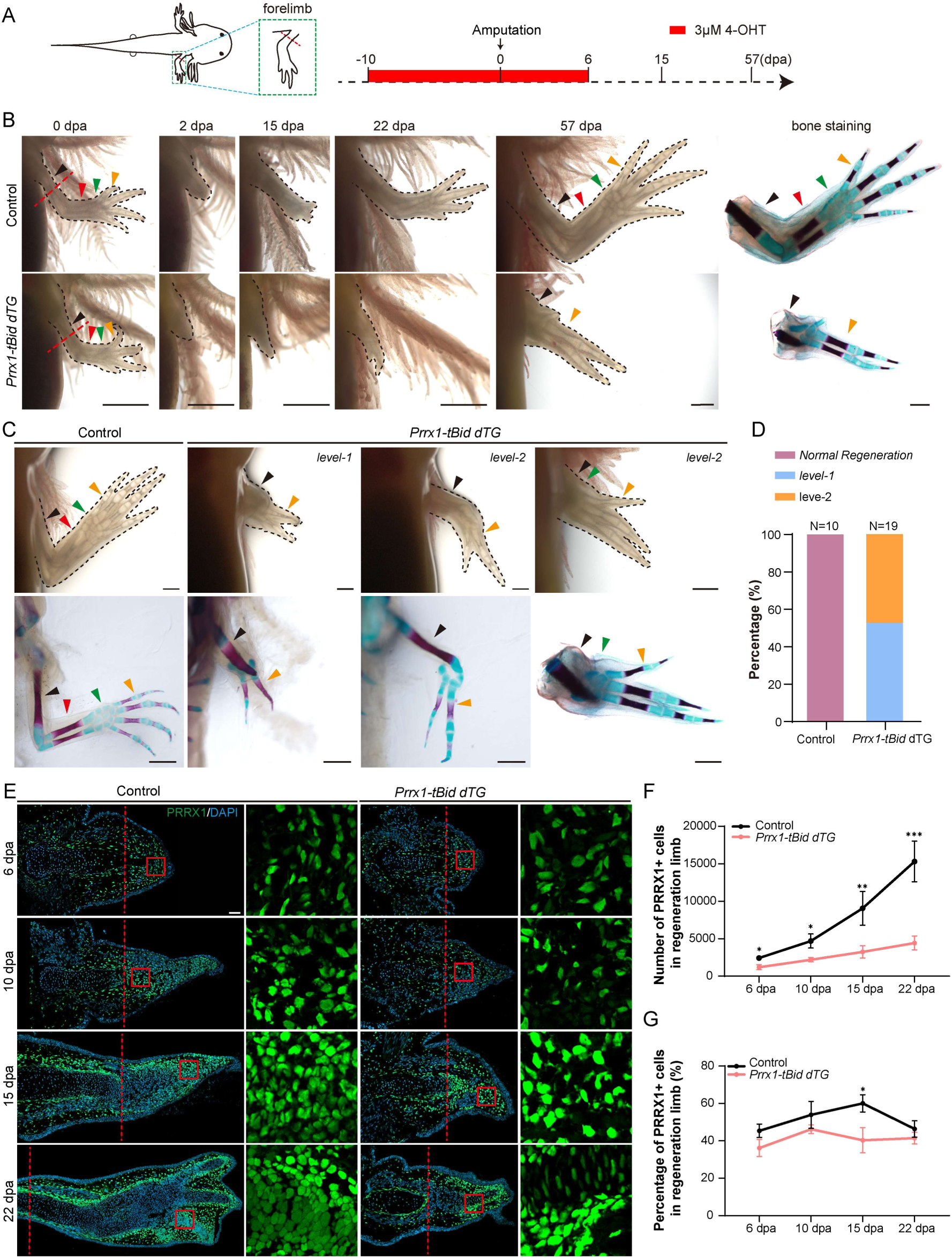
Early ablation of CT cells impairs the regeneration of both upper and lower limbs, while having a lesser effect on hand outgrowth. A) Timeline of 4-OHT treatment during early regeneration in *Prrx1-tBid dTG* axolotls. B) Representative time-series images of limb regeneration after 4-OHT treatment in control (upper panel) and *Prrx1-tBid dTG* (lower panel) axolotls. The right column shows the skeletal elements at 57 days post-amputation (dpa) via alcian blue/alizarin skeleton staining. The limb regeneration defected in *Prrx1-tBid dTG* axolotls compared to controls. C) Live imaging (upper panel) and Alcian blue/Alizarin red skeletal staining (lower panel) of regenerated forelimbs from *Prrx1-tBid dTG* and control axolotls at 57 dpa, following 4-OHT treatment. Two distinct regenerated phenotypes were observed following CT cell ablation. D) Phenotypic analysis of regenerated forelimbs based on Alcian blue/Alizarin red skeletal staining. E) Immunofluorescence of PRRX1 (green) and DAPI (blue) on limb longitudinal-sections from control and *Prrx1-tBid dTG* axolotls at 6, 10, 15 and 22 dpa. The *Prrx1-tBid dTG* axolotls (right) show a significant reduction in PRRX1+ cells compared to controls (left). Red boxes outline the higher magnification area of limb bud, showing single-channel. F) and G) Quantification of the absolute number (F) and percentage (G) of PRRX1+ cells within the newly formed blastema and a combined 500 μm zone proximal to the amputation plane at 6, 10, 15 and 22 dpa in control (black line, n = 3) and *Prrx1-tBid dTG* (red line, n = 3) axolotls. Red dashed lines indicate amputation planes. Black dashed lines outline the shape of limb. Black arrowheads indicate the stylopods; red arrowheads indicate the zeugopods; green arrowheads indicate the autopods; yellow arrowheads indicate the digits. For quantification, data were analyzed by unpaired two-tailed Student’s t-test and represented as mean ± SEM, \**p* <0.05; \*\**p* <0.01, \*\*\**p* <0.001. Scale bars: 1 mm in (B) and (C); 100 μm in (E).

We next analyzed the phenotypes upon CT ablation at cellular level during limb regeneration. Immunofluorescence staining of longitudinal sections across regeneration stages (6, 10, 15, and 22 dpa) showed that *Prrx1-tBid dTG* regenerates were smaller, exhibited delayed distal structure formation, and contained markedly fewer PRRX1+ cells compared to controls. By 22 dpa, limb architecture in *Prrx1-tBid dTG* axolotls appeared markedly disorganized, showing the most pronounced structural distinction from controls (Figure 3E). Quantitative analysis further confirmed that although PRRX1+ cell numbers increased progressively in both groups, the counts remained consistently lower in *Prrx1-tBid dTG* limbs across all stages (Figure 3F). The difference in PRRX1+ cell numbers between the two groups widened over time and reached a plateau by 22 dpa. Moreover, the proportion of PRRX1+ cells within the blastema was significantly reduced in ablated animals, with a particularly notable decrease observed at 15 dpa (Figure 3G). The regenerative phenotypes after CT cell ablation to some extent recapitulated those previously observed in early development. Collectively, these results demonstrate that CT cells are essential for successful limb regeneration in axolotls. And the phenotypic parallels observed following CT ablation point to a potential common regulatory mechanism mediated by CT cells in both limb development and regeneration.

### CT ablation results in reduction of the upper and lower limb domains during limb regeneration and development

The expression of *Hoxa9*, *Hoxa11*, and *Hoxa13* is known to be spatiotemporally correlated with limb segment specification during proximal-to-distal limb development and regeneration, providing a molecular framework for segmental patterning and identity^[38, 54]^. To determine whether CT ablation disrupts this *Hoxa* expression profile and early limb domain patterning, we analyzed the expression of *Hoxa9, Hoxa11,* and *Hoxa13* in regenerating forelimb blastemas using mRNA in situ hybridization. None of the three *Hoxa* genes were detectable in blastemal cells from either control or *Prrx1-tBid dTG* axolotls at 2dpa. At 4 dpa, the expression of *Hoxa9* and *Hoxa11*, but not *Hoxa13*, became evident and spread throughout the blastemas in both groups. By 6 dpa, a nested expression pattern of these three genes was observed in the blastemas of controls, clearly demarcating three distinct limb domains^[38]^. In contrast, *Prrx1-tBid dTG* blastemas exhibited a pronounced reduction in the size of both the upper (*Hoxa9+*) and lower (*Hoxa9+/ Hoxa11+*) limb domains, whereas the *Hoxa13*-expressing hand domain was less affected compared to controls (Figure 4A,C). A parallel analysis during limb development showed similar alterations in *Hoxa* gene expression in *Prrx1–tBid dTG* limb buds at 11 dpt (Figure 4B,D). These findings align with the segmental defects observed following CT ablation (Figure 2D and 3B). Notably, previous work demonstrated that misexpression of *Hoxa13* in the proximal wing bud reduces zeugopod size in chick ^[76]^, suggesting that the relative expansion of the *Hoxa13* domain versus the *Hoxa9*/*Hoxa11* domains could contribute to proximal segment reduction during regeneration. Together, these data indicate that early CT ablation leads to a pronounced decrease in the *Hoxa9* and *Hoxa11* expression domains—corresponding to the upper and lower arm segments—while sparing the distal hand domain during both development and regeneration. This shift ultimately results in loss of the zeugopod but preservation of the autopod.

**Figure 4.**
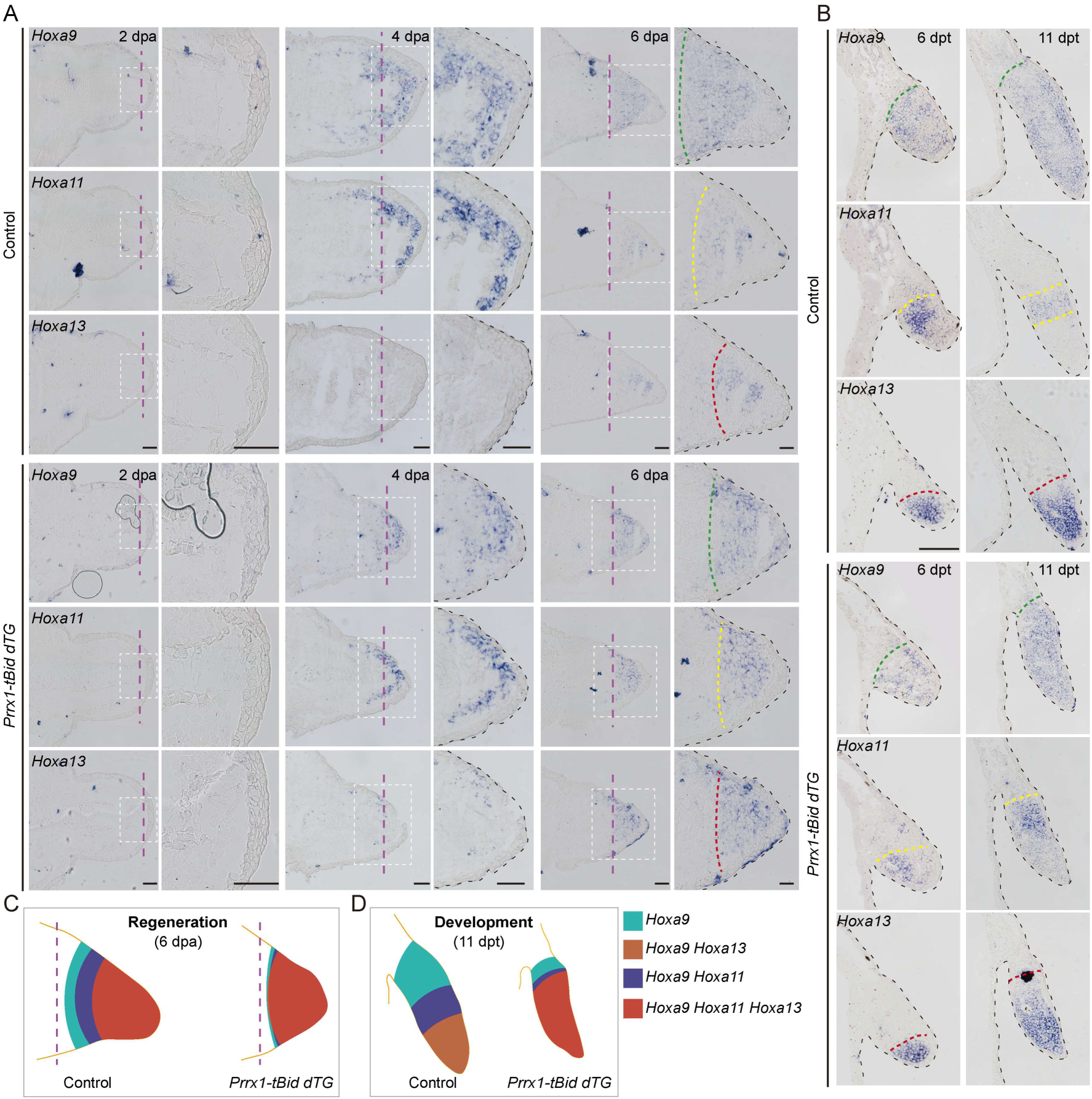
CT cell ablation leads to reduction of the proximal but not the distal limb domains. A) and B) Representative time-series images of in situ hybridization (ISH) for *Hoxa9* (upper panels), *Hoxa11* (middle panels) and *Hoxa13* (lower panels) on adjacent longitudinal-sections of blastemas during regeneration (A) or limb buds during development (B) from 4-OHT treated control (*Caggs^EGFP/tBid^*) and *Prrx1-tBid dTG* axolotls. Purple dashed lines in (A) indicate amputation planes. White boxes in (A) outline the higher magnification area of limb. Black dashed lines outline the shape of limb blastemas (A) and limb buds (B). Green, yellow, and red dashed lines outline the expression boundaries of *Hoxa9*, *Hoxa11*, and *Hoxa13*, respectively. C) and D) The cartoon schematic shows the expression of *Hoxa9*, *Hoxa11* and *Hoxa13* in blastemas (C) and limb buds (D) of control (left) and *Prrx1-tBid dTG* (right) axolotls. Purple dashed lines in (C) indicate amputation planes. Turquoise area represents *Hoxa9* expression domain; navy blue area, overlapping *Hoxa9* and *Hoxa11* domains; brown area, overlapping *Hoxa9* and *Hoxa13* domains; red area, overlapping *Hoxa9*, *Hoxa11*, and *Hoxa13* domains. Note that the decrease in size of the domains (*Hoxa9*) and (*Hoxa9*, *Hoxa11*) corresponding to the upper and lower limbs upon CT ablation, compared to the controls. Scale bars: 100 μm.

### Mapping CT subtypes and their dynamics during limb regeneration

To characterize the molecular signature and dynamics of CT cells during limb regeneration, we constructed a comprehensive single cell atlas of limb regeneration using the control axolotls. We adopted a droplet-based scRNA-seq (SeekOneDigital Droplet System), and collected samples from four key time points for analysis, including pre-amputation (0 dpa, upper limb region), 6, 10 and 15 dpa. After applying quality controls^[77, 78]^, 43008 high-quality cells were retained, characterized by a median of 3078 genes per cell (Figure 5A and Figure S6A,B, Supporting Information). Following clustering identification in a low-dimensional embedding^[77]^, biological replicates exhibited high similarity in this embedding (Figure S6C, Supporting Information) and an even distribution across clusters. Based on the expression of marker genes, single-cell transcriptomic profiling of the *Caggs^EGFP/tBid^* axolotl limb regeneration identified 17 distinct cell populations which cover all major limb cell types^[7, 24, 79, 80]^ (Figure S6D, Supporting Information). It included CT cell (*Prrx1*), chondrocyte (*Sox9*, *Acan* and *Matn4*), tenocyte (*Aspn* and *Timp1*), skeletal muscle cell (*Myf5*, *Mylpf* and *Ckm*), MuSC (*Pax7*), keratinocyte (*Krt5*), apical epithelial cap (AEC) cell (*Col17a1* and *Vwa2*), erythrocyte cell (*Hbe1* and *Hmbs*), neuron (*Insm1* and *Scng*), schwann cell (*Mpz* and *Foxd3*), epidermal cell (*Krt12* and *Epcam*), vascular endothelial cell (*Vwf*, *Plvap* and *Pecam1*). Additionally, we also detected immune cells, such as macrophage (*C1qb, C1qa, Marco* and *Cd68)*, cytotoxic T cell (*Cd3e*, *Cd3g* and *Gzma*), mast cell (*Kit* and *Cpa3*), and neutrophil-like cell (*Csf3r* and *Mmp9*) (Figure 5B and Figure S6E, Supporting Information).

**Figure 5.**
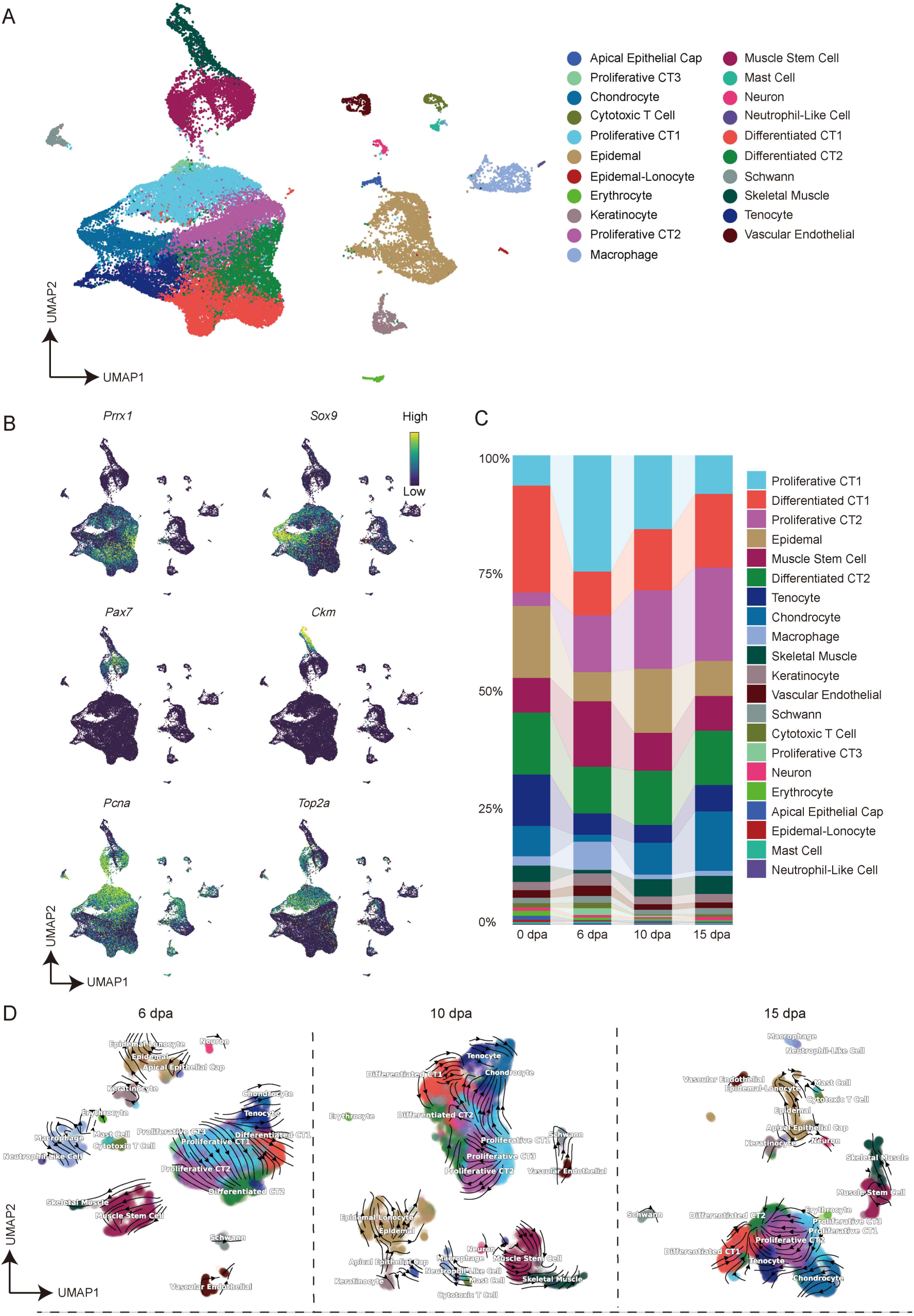
scRNA-seq reveals proximal-distal transition of nascent CTs during limb regeneration. A) UMAP plot showed clustering of 43008 cells into 21 distinct populations in control axolotl limb regeneration. B) UMAP plot revealed distinct cellular territories defined by key markers: CT (*Prrx1*), chondrocyte (*Sox9*), muscle stem cells (*Pax7*), skeletal muscle (*Ckm*), and proliferation markers (*Pcna*, *Top2a*) in regenerating axolotl limbs. C) Stacked histograms depict sample-wise cell type distributions, expressed as proportions relative to individual sample cellularity. D) RNA velocity streamline plots visualize the predicted trajectory of cell lineage transitions at 6, 10 and 15 dpa in regenerating control axolotl limbs, with cells are colored according to annotated cell types.

The CT was characterized primarily by the expression of *Prrx1* (Figure 5B and Figure S6E, Supporting Information), which was subjected to a rapid expansion post-injury (Figure S6F, Supporting Information). We next subclustered the CT population and analyzed their dynamics during regeneration. We stratified all undefined CT populations into five distinct sub-populations, including proliferative CT1, proliferative CT2, proliferative CT3, differentiated CT1 and differentiated CT2, and investigated their dynamics during regeneration. Sub-populations proliferative CT1 and proliferative CT2 cells, were identified and marked by the expression of proliferation markers *Pcna* and *Top2a* (Figure 5A,B and Figure S6E,G, Supporting Information), suggesting their progenitor property. In response to limb injury, the proliferative CT1 was rapidly expanded, peaked at early stage of injury (6 dpa) in regenerating axolotl limbs, and subsequently decreased, similar to the dynamics observed in macrophage cells at the early healing process^[22, 25, 27]^ (Figure 5C). In contrast, proliferative CT2 sustained a progressive accumulation following their initial rise (Figure 5C). The other two sub-populations, differentiated CT1 and differentiated CT2 were characterized by high expression of *Lum* and *Dpt*, and *Tbx15* respectively^[24, 37]^ (Figure S6E, Supporting Information). However, the proportion of differentiated CT1 cells displayed dynamic changes after injury, with an initial decrease followed by a gradual increase, in contrast to the fairly constant ratio maintained by differentiated CT2 during regeneration (Figure 5C). The proliferative CT3 sub-cluster only occupied a small proportion in the entire undefined CT population (Figure 5A,C). Further RNA velocity analysis^[24, 81]^ revealed that the proliferative CT1 subtype is the cell source, the primary respondent to injury, giving rise to proliferative CT2 and subsequently differentiating into differentiated CT1/CT2 cells (Figure 5D). Collectively, our data delineate the injury-responsive CT subtypes and their lineage trajectory in generating proximal CT cells.

### Early CT ablation causes differentiated CT1 rapid decline and delayed distal transition of subpopulation of CT progenitors

The forelimb regenerates in *Prrx1-tBid dTG* axolotls displayed severe defects after early CT ablation, characterized by the loss of zeugopod and/or autopod between the upper arm and digits (Figure 3B,C). To investigate the underlying CT-mediated regulatory mechanisms responsible for forelimb regeneration defects, we performed scRNA-seq on limb samples collected at 6, 10, and 15 dpa from CT-ablated *Prrx1-tBid dTG* axolotls. Following quality control, we integrated the single-cell transcriptomic datasets from both *Prrx1-tBid dTG* and control axolotls, yielding 75904 high-quality single-cell transcriptomic profiles. We next annotated these data and identified 21 cell types in total using the unsupervised clustering analysis based on gene expression (Figure 6A and Figure S7A,B, Supporting Information). Biological replicates showed consistent representation across all annotated cell populations (Figure S7C, Supporting Information).

**Figure 6.**
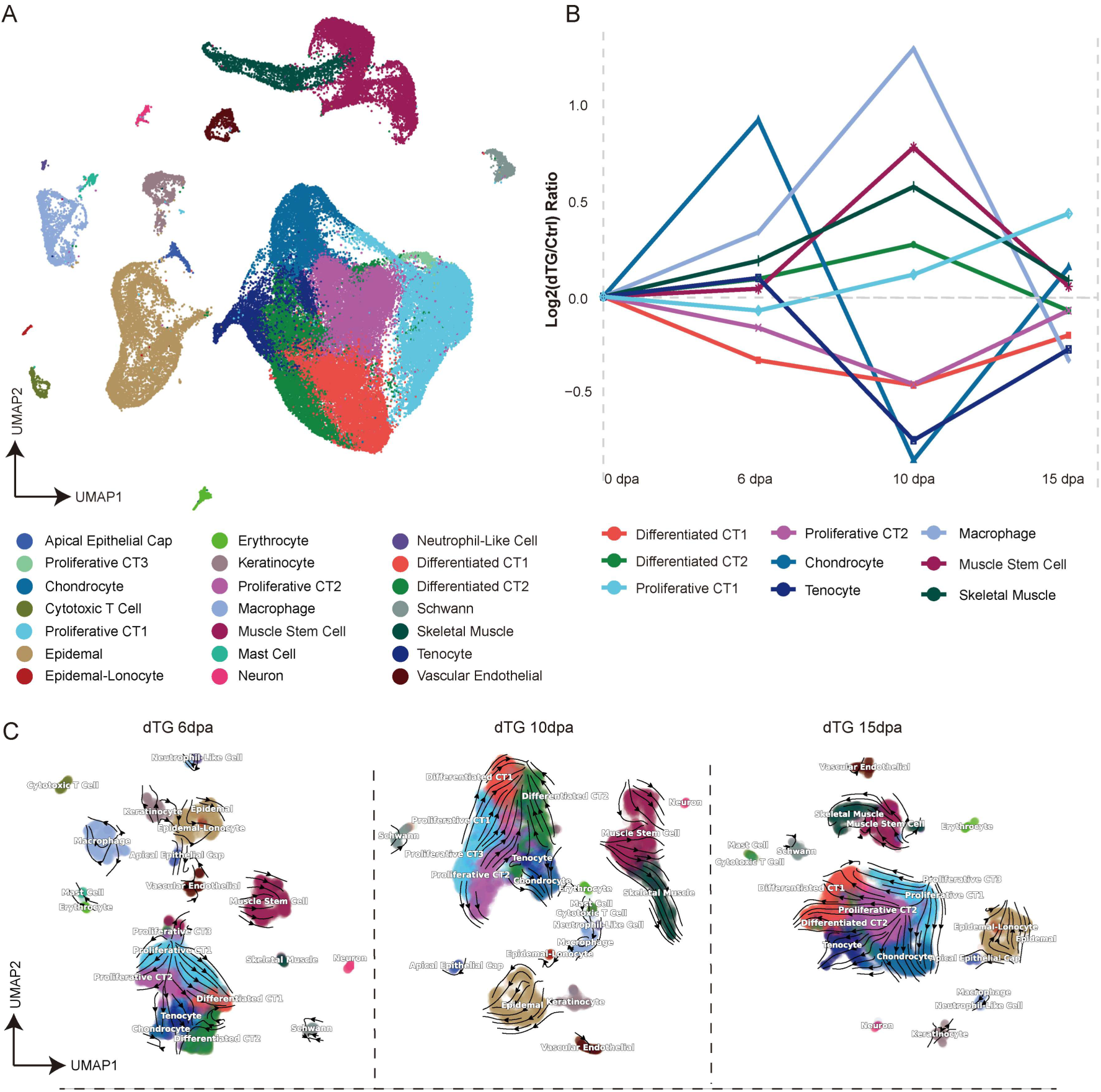
scRNA-seq reveals accelerated distal migration within the CT subpopulations of *Prrx1-tBid dTG* axolotls compared to controls. A) UMAP plot showed clustering of 75904 cells into 21 distinct populations in *Prrx1-tBid dTG* and control (Ctrl) axolotl limb regeneration. B) Line graph showing the log_2_ cell proportion ratios (*Prrx1-tBid dTG* axolotls vs. controls) across all stages. Values >0 indicate higher cell proportions in *Prrx1-tBid dTG* axolotls relative to controls, while values <0 reflect the inverse relationship. C) RNA velocity streamline plots depict the predicted trajectory of cell lineage transitions across key regenerative time points (6, 10 and 15 dpa) in *Prrx1-tBid dTG* axolotls limb regeneration. Each cell is colored by its annotated cell type.

To investigate the cellular interference causing the forelimb regeneration defects, we explored the dynamics of nine cell types, including the major CT subtypes by quantifying the proportional changes of each population in *Prrx1-tBid dTG* axolotls versus controls across three regenerative time points (Figure 6B and Figure S8A, Supporting Information). Notably, differentiated CT1 declined massively in *Prrx1-tBid dTG* axolotls at early regeneration stage (6 dpa) and exhibited persistently lower proportions throughout the regeneration process, compared to controls. This differential was progressively diminished after 10 dpa, suggesting regenerative compensation of differentiated CT1 following the loss of PRXX1+ cells (Figure 6B and Figure S8A, Supporting Information). Differentiated CT1 cells retained their proximal identity throughout regeneration, as evidenced by sustained high expression of the proximal markers *Meis2* and *Shox* (Figure S8B,C, Supporting Information). Given the observed loss of the upper arm phenotype upon CT ablation, it indicates that the differentiated CT1 is the predominantly affected subpopulation in CT-ablated *Prrx1-tBid dTG* axolotls.

We next compared the regeneration dynamics of proliferative CT1 and proliferative CT2 progenitor cells, which are two progenitor subtypes critical for CT regeneration. At 10 and 15 dpa, we observed a significantly higher proportion of proliferative CT1 cells and a concomitantly lower proportion of proliferative CT2 cells in *Prrx1-tBid dTG* limbs compared to controls, despite comparable levels at 6 dpa (Figure 6B and Figure S8A, Supporting Information). However, RNA velocity analysis revealed that the limb regeneration after CT ablation in *Prrx1-tBid dTG* axolotls followed a trajectory similar to that in axolotls^[24, 81]^ (Figure 6C). These observations revealed that the progression, but not the lineage from the proliferative CT1 to proliferative CT2 was likely delayed in *Prrx1-tBid dTG* axolotl.

To explore the mechanisms underlying the altered CT progenitor progression after CT ablation in *Prrx1-tBid dTG* axolotls, we first characterized dynamics and transition in the identity of CT subtypes during limb regeneration. We analyzed the expression of *Hoxa13*, *Hoxd13*, *Meis2*, and *Shox*^[49, 52, 58]^, which are known to play roles in establishing proximal and distal axis by providing positional identity during limb regeneration. The results showed that, in both the control and *Prrx1-tBid dTG* axolotls, the distal markers *Hoxa13*, *Hoxd13* were enriched in proliferative CT1 and proliferative CT2. These two markers exhibited low expression at the early stage of regeneration (6 dpa), and high expression at the middle and late stages (10 and 15 dpa). Conversely, the proximal markers, *Meis2* and *Shox*, were predominantly enriched in the CT subtypes differentiated CT1 and differentiated CT2, showing high expression at the early and middle stages of regeneration (6 and 10 dpa), but low expression at the late stage (15 dpa) (Figure S9A,B, Supporting Information). Although applying this small marker set can show a transformation trend of CT subpopulations from proximal to distal, it fails to distinguish the difference between these two groups.

Next, we composed proximal and distal gene modules for defining CT identity using published distal-proximal differentially expressed genes, including genes such as *Hoxa13*, *Hoxd13*, *Meis2*, and *Shox* (Table S1, Supporting Information)^[52]^. We analyzed the distal and proximal gene module activity in CT subpopulations during limb regeneration. Strikingly, in both control and *Prrx1-tBid dTG* axolotls, the global pattern of CTs exhibited a strong trend of progressive transition of the positional information, first weakening the proximal and then strengthening the distal identity following the process of limb regeneration (Figure 7A and Figure S10A,B, Supporting Information). The sub-populations of proliferative CT1, and proliferative CT2 displayed enriched proximal gene module activity at the early regeneration phase (6 dpa), but shifted to higher distal scores at later stages (10 and 15 dpa) in both groups (Figure 7A and Figure S10A,B, Supporting Information). Together with the quantitative dynamics and trajectory analysis of CT cells (Figure 5C,D, Figure 6B,C and Figure S9A, Supporting Information), it was identified that the proliferative CT1 are major CT cells and the potential proximal progenitor sources contributing to distal transformation and generating distal CT cell population during limb regeneration. In contrast, differentiated CT1 constantly exhibited high proximal gene module activity throughout regeneration, indicating differentiated CT1 is one of the perspective residential CT subtypes maintaining their proximal identity during regeneration in both control and *Prrx1-tBid dTG* axolotls (Figure 7A and Figure S10A,B, Supporting Information). Such pattern of temporal shift of CT subtype positional identity was consistent with theoretical proximal-to-distal transition in limb regeneration. And the overall tendency of each CT subtype showed similar proximal/distal dynamics during regeneration in both control and *Prrx1-tBid dTG* axolotls.

**Figure 7.**
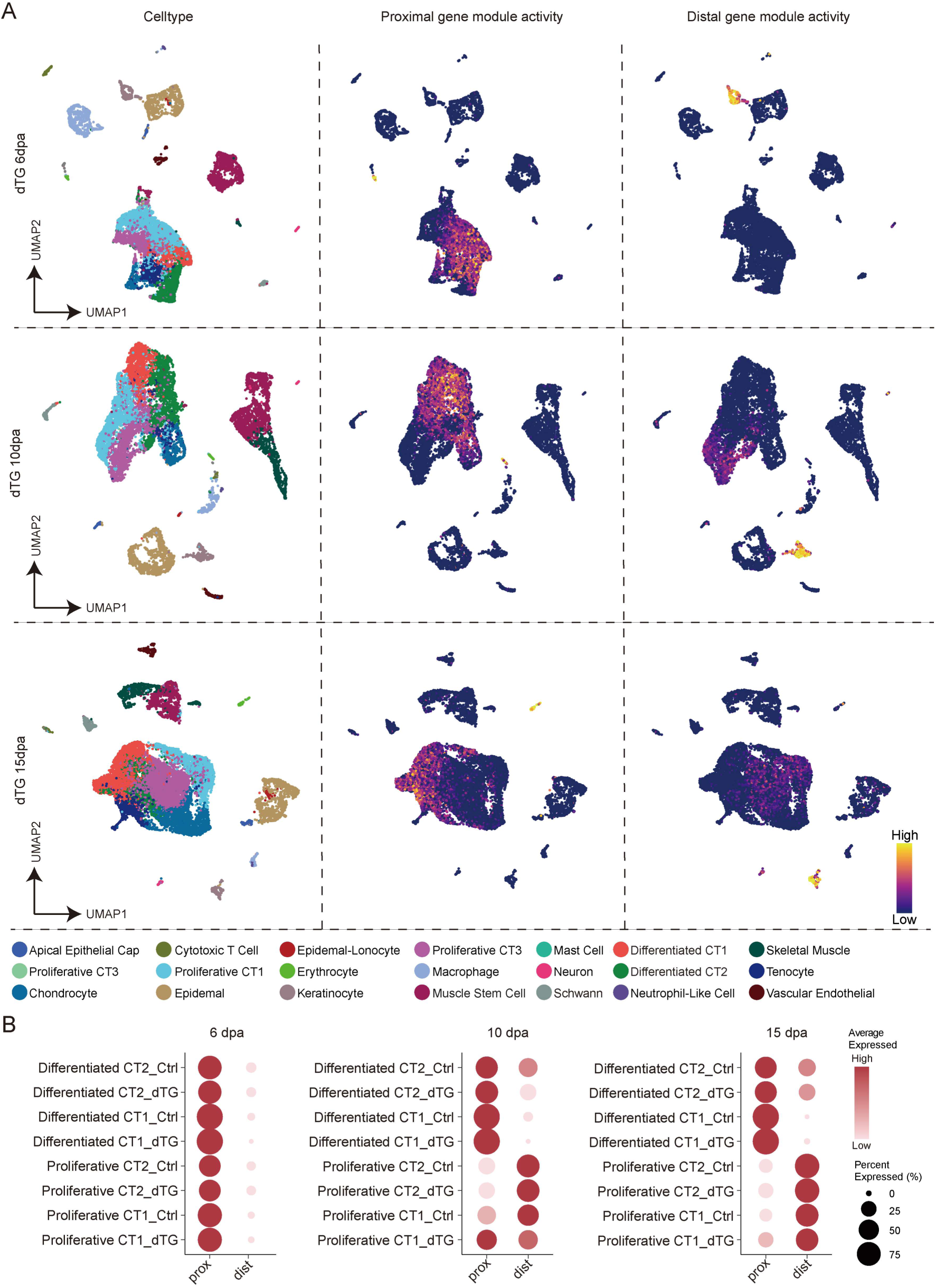
Analysis of gene module activity in limb distalization. A) UMAP depicting the enrichment of distal and proximal gene modules in control (Ctrl) limbs. B) The dot plot demonstrates enrichment of gene module activity in distal (dist) and proximal (prox) regions of *Prrx1-tBid dTG* and control axolotls across limb regeneration.

We then investigated the proximal-distal transition after CT ablation. We identified that, at 10 dpa, proliferative CT1 cells in *Prrx1-tBid dTG* axolotls exhibited a higher proximal module activity, yet lower distal module activity compared to the controls (Figure 7B). It suggested a delayed distal transition in proliferative CT1 after CT ablation, consistent with the delayed limb regeneration phenotype observed in the *Prrx1-tBid dTG* axolotls after CT ablation (Figure 3B and Figure S5A, Supporting Information). In summary, based on the analysis using composed proximal and distal gene modules, we identify major CT subtypes respectively contributing to proximal and distal regeneration, and a progressive proximal-to-distal transition in CT cells during limb regeneration. Moreover, CT ablation causes a delayed distal transformation of the proliferative CT1, one of the major CT subtypes. It may consequently lead to an overall retardation of the limb regeneration compared to the controls.

### CT cells regulate adaptive proliferation of muscle stem cells following CT ablation

In response to CT ablation, the entire defined limb segment (e.g., lower limb) was absent. Not only the ablated CT tissues, but also other surrounding tissues, such as muscle lineage adjusted their behavior to adapt to the situation of CT loss to produce a smaller number of muscles. It implied the dominant role of CT and their interaction with other adjacent tissues during regeneration. To investigate how muscle tissue respond to CT ablation to adjust their proliferative behavior, we performed limb amputations and collected samples from the *Prrx1-tBid dTG* and control axolotls as described above, with an additional single EdU pulse prior to sample collection, to analyze the dynamics of MuSCs during regeneration. Immunohistochemistry on longitudinal sections using antibodies against PAX7 (MuSCs) and PRRX1 (CT), combined with EdU detection, showed that the total number of PAX7+ MuSCs increased over time in both groups, but was significantly reduced in *Prrx1-tBid dTG* axolotls at 15 and 22 dpa compared with corresponding controls (Figure 8A,B). Consistently, quantification of EdU+/PAX7+ cells or analysis of the cycling cells from scRNA-seq data revealed that proliferating MuSCs exhibited similar dynamics as the total PAX7+ cells during regeneration in both groups (Figure S11A-D, Supporting Information). Following the trend of declined total number and proliferating MuSCs along regeneration, conversely, the ratio of PAX7+ cells to PRRX1+ cells were markedly elevated in *Prrx1-tBid dTG* axolotls, gradually increased at 10, 15, then declined at 22 dpa, compared to controls (Figure 8C). These data indicate that, following sudden PRRX1+ CT ablation in the *Prrx1-tBid dTG* axolotls, PAX7+ cells gradually reduce their proliferation to adapt to the CT decrease, and eventually reach a balance. It further suggests that CT communicate with surrounding cells, including MuSCs to regulate their proliferation for proper size control during limb regeneration.

**Figure 8.**
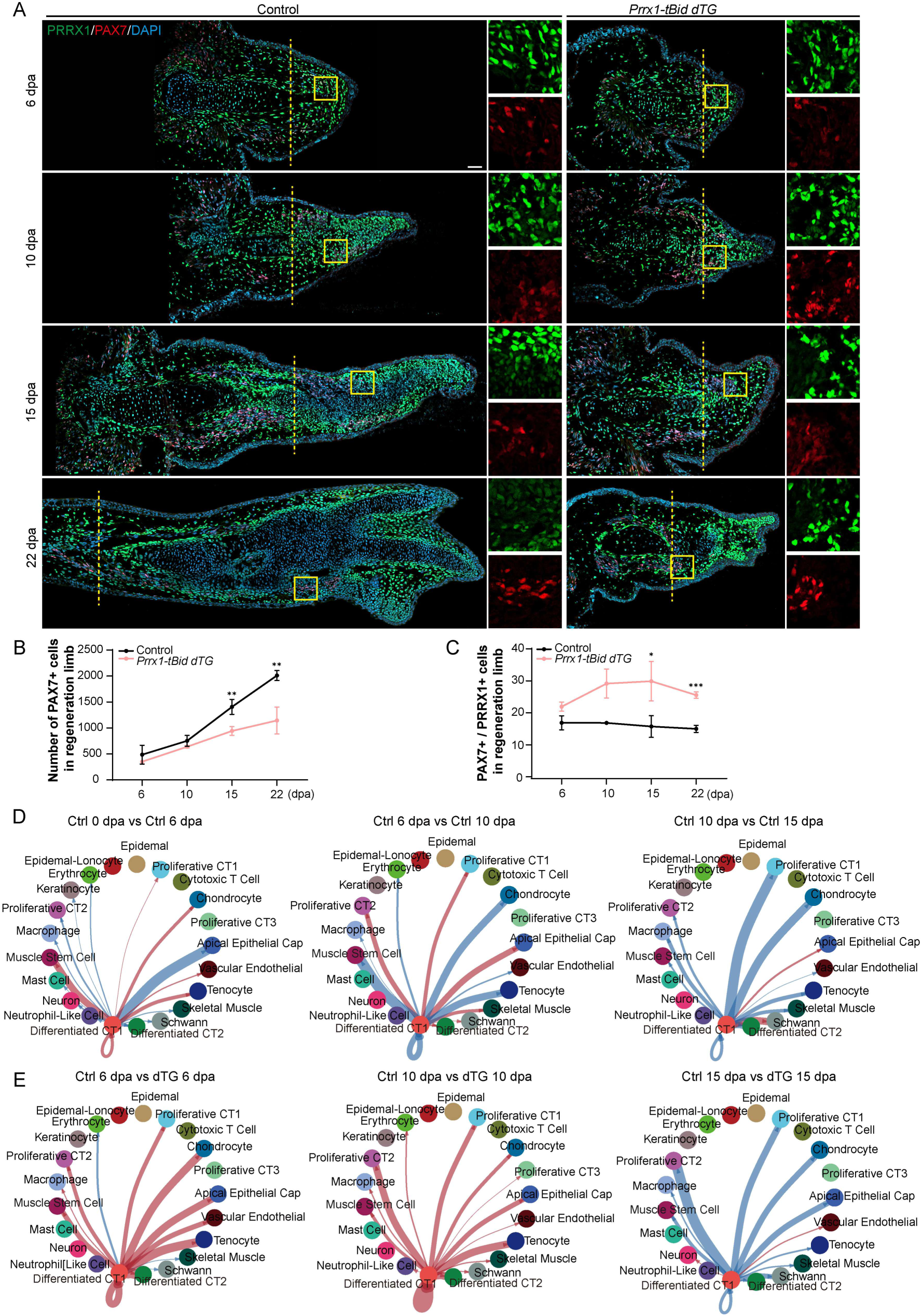
CT cell ablation perturbs muscle stem cell dynamics and intercellular crosstalk. A) Immunofluorescence images of PRRX1 (green), PAX7 (red), and DAPI (blue) on limb longitudinal-sections from control (left) and *Prrx1-tBid dTG* (right) axolotls at 6, 10, 15 and 22 dpa. Yellow dashed lines indicate amputation planes. Yellow boxes outline the higher magnification area of blastema, showing single-channel. B) Quantification of the absolute number of PAX7+ cells within the newly formed blastema and a combined 500 μm zone proximal to the amputation plane at 6, 10, 15 and 22 dpa in control (black line, n = 3) and *Prrx1-tBid dTG* (red line, n = 3) axolotls. C) Quantification of the percentage of PAX7+ cells among the PRRX1+ cell population within the same region described in (B) for control (black line, n = 3) and *Prrx1-tBid dTG* (red line, n = 3) axolotls. D) Chord diagrams depict cell-cell communication patterns between differentiated CT1 and other cell types in control samples at sequential regeneration stages: 0 vs 6 dpa (left), 6 vs 10 dpa (middle), and 10 vs 15 dpa (right). Red connections indicate stronger interactions in the latter time point relative to the former, whereas blue denotes weaker interactions in the latter compared to the former. E) Chord diagrams compare differentiated CT1 and other cell types interaction networks between *Prrx1-tBid dTG* and control axolotls at 6, 10, and 15 dpa. Red connections indicate stronger interactions in *Prrx1-tBid dTG* axolotls relative to controls, whereas blue denotes weaker interactions in *Prrx1-tBid dTG* axolotls versus controls. For quantification, data were analyzed by unpaired two-tailed Student’s t-test and represented as mean ± SEM, \**p*<0.05; \*\**p*<0.01; \*\*\**p*<0.001. Scale bar: 100 μm.

Cellular crosstalk in the local microenvironment orchestrates the process of tissue regeneration^[82]^. To explore the potential cell-cell communications between CT and other cell types during regeneration, we next analyzed the interaction profiles of each cell type in the control axolotls (Figure S12A, Supporting Information). Given that differentiated CT1 represents the primary CT subpopulation ablated in *Prrx1-tBid dTG* animals, we focused specifically on the interactions between differentiated CT1 and other cell types. We found that differentiated CT1 predominantly communicate with proliferative CT1, proliferative CT2, chondrocytes, the apical epithelial cap cells, tenocytes, and MuSCs (Figure S12B, Supporting Information). To better understand the dynamics of cell-cell interactions during limb regeneration in control axolotls, we performed pairwise comparisons of interaction strengths across different regenerative timepoints. Our analysis revealed that the interaction between differentiated CT1 and MuSCs was stronger at 6 dpa than at 0 dpa, weaker at 10 dpa compared to 6 dpa, and absent by 15 dpa (Figure 8D). Notably, a global downregulation of intercellular interactions was observed at 15 dpa compared to 10 dpa (Figure 8D), possibly reflecting a transition toward cellular homeostasis in the late regenerative stage, during which cells may exhibit reduced communicative activity. Subsequently, we investigated the cell-cell interactions in *Prrx1-tBid dTG* after CT ablation and analyzed the variations between CT-ablated *Prrx1-tBid dTG* and control during regeneration. The results revealed enhanced interaction strengths in *Prrx1-tBid dTG* axolotls compared to controls at 6 and 10 dpa, particularly in cell types such as tenocyte, chondrocyte and MuSC (Figure 8E and Figure S13, Supporting Information). In contrast, by the late regenerative stage (15 dpa), a marked downregulation of intercellular communication was observed in *Prrx1-tBid dTG* axolotls relative to controls (Figure 8E). These data showed that CT ablation induced an elevated and prolonged communications between CT and adjacent MuSCs at early regeneration stages, probably to modulate the adaptive proliferation of MuSCs to the CT ablation. The level of cell-cell communications reached the peak at early stage of regeneration and gradually returned to level of uninjured status. Overall, these data indicate that CT is one of the major cell types interacting with other surrounding cells during limb regeneration.

## Discussion

Our findings established CT as a master regulator of axolotl limb patterning and growth. Using a *tBid*-mediated ablation system, we showed that CT provides essential positional information, and its temporal attenuation led to specific limb segment loss. scRNA-seq revealed that CT subtypes undergo a proximal-to-distal transformation, and their ablation resulted in the reduction and proper proximal/distal transition disruption of certain CT subclusters. Moreover, CT orchestrated the behavior of surrounding tissues, positioning it as a central coordinator of limb regeneration.

The *tBid*-mediated conditioned targeted cell ablation method provides a powerful tool for studying the function of specific cell type and the interactions between cell types during developmental or physiological processes. Currently, there are several methods designed for genetic ablation of targeted cell types in various organisms, such as diphtheria toxin A, Kid/Kis, and NTR-Mtz. Tissue-specific expression of diphtheria toxin A in mice and zebrafish can lead to effective ablation of target cell populations, but it may also lead to non-specific death of other cells or even death of organism^[83, 84]^. The Kid/Kis system is also used to achieve cell ablation in zebrafish, which is to express the bacterial toxin Kid in the target population while expressing its antidote Kis in the remaining cells to protect these cells from KID-induced cell death, but this method can only be used in embryos with instantaneous Kid/Kis expression^[67, 68]^. NTR represents a potential cell ablation method for salamanders and has been primarily applied in zebrafish. NTR catalyzes substrates such as MTZ to yield toxic products to cause specific cell ablation^[61–64]^. Two recent studies showed that NTR/Mtz system achieves ablation of CTs, neural stem cells and neurons in the central nervous system^[26,^

^85]^. There are generally two strategies to reach targeted cell ablation using NTR/Mtz system, to directly control NTR expression with tissue specific promoters, or to combine an inducible Cre line in which Cre expression is driven by tissue-specific promoters and a “floxed” NTR line. The former application requires developing specific transgenic line. In contrast, the latter approach could employ any available inducible Cre line to achieve tissue-specific NTR expression. In this case, two drugs, e.g., Tamoxifen and Mtz need to be applied to animals. In our study, we chose *tBid* as the trigger of cell ablation. *tBid* is a pro-apoptotic factor of endogenous apoptosis pathway which induces apoptosis in target cells without causing an inflammatory response, and therefore minimizes the potential damage to surrounding cells. And when combining with inducible Cre lines to achieve tissue-specific tBid expression and cell ablation, administration of a single drug tamoxifen is sufficient.

The limb is a complex structure comprised of varied types of tissue. It has been shown that tissues such as muscle lineage and bone are dispensable for determining the positional identity during limb regeneration^[14, 25, 28, 30, 86, 87]^. In contrast, CTs produce many types of progenies during limb development and regeneration and actively interact with surrounding tissues^[24, 26, 28–31]^. It was indicated that CTs play critical roles in regulating the positional information during limb development and regeneration^[14, 30, 35, 37, 48–50]^. Our results demonstrated that genetic ablation of CTs caused loss of limb segments during development and regeneration. When ablating CTs at early developmental or regeneration stages, a portion of limb segments in between shoulder and hand were missing, but the most distal hand was less affected. It provided direct evidence to support the essential role of CTs in limb development and regeneration.

During limb development and regeneration, CTs from different limb segments express different markers that in turn determine the positional identity of limb segments. *Hoxa* genes are one of such markers predominantly expressed in the PRRX1+ CT cells^[38]^. The progressive onset expression of *Hoxa9*, *Hoxa11* and *Hoxa13* in CTs of developing limb bud follows a chronological order, associated with the formation from proximal to distal limb segments^[38]^. The *Hoxa* gene expression in blastema during limb regeneration exhibits a similar pattern as observed during development. From the lineage perspective, residential CTs in distal segments (e.g., hand) should be the descendant derived from CTs in proximal segments (e.g., upper limb). The lineage relationship is particularly obvious during regeneration upon amputation at the position of upper limb^[24, 28, 30]^. While ablating CTs at early development or regeneration, we would assume that CT cells expressing *Hoxa9+* and/or *Hoxa9+/Hoxa11+*, corresponding to proximal limb segments were primarily eliminated in limb bud/blastema. In situ hybridization indeed showed a dramatic reduction of the upper-limb (*Hoxa9*+) and lower-limb (*Hoxa9*+/*Hoxa11*+) domains, but not the hand domain (*Hoxa9*+/*Hoxa11*+/*Hoxa13*+), at early developmental and regenerative stages after CT ablation.

After early CT ablation followed by upper limb amputation, the remaining residential CT cells in proximal limb must be the source contributing to the distal limb segment regeneration. It meant that these proximal residual CT cells neither maintain their original identity to grow or regenerate a normal limb in a temporal delayed manner, nor give rise to a smaller upper limb on time. The phenotypic observations suggested that these CT cells proliferate and commit to more distal identity, involved in distal structure (e.g. hand) formation/regeneration. It is consistent with the in-situ hybridization revealing that the relative expansion of *Hoxa13*-domain and reduction of non-*Hoxa13*-domain compared to that in the controls at 6 days post amputation. scRNA-seq analysis further showed that upon CT ablation, some CT cells tended to be delayed on distal transformation, explaining the observed delayed digit formation phenotype during limb regeneration after CT ablation (Figure 7B). Collectively, these results suggested that the transformation of CT from proximal to distal identity during limb development or regeneration follows a continuous and strict temporal order. Once interrupted, proximal-distal transformation may proceed in a discontinuous manner, following a predefined chronological order with the interrupted portion being disregarded (<u>loss of relevant segment missing the critical time</u>).

Moreover, our data showed that in response to the CT ablation, all other surrounding tissues such as muscles eventually adapted proportionally to the reduction of CTs, leading to the loss of the entire limb segments, including the progeny derived from CT and other progenitors. We found that upon CT ablation, MuSCs expanded in a similar manner as that in the controls, yielding similar number, but a higher ratio of MuSCs at early regeneration stage. In contrast, at late regeneration phase, MuSCs decrease their proliferating rate/speed to adapt to the loss of CT cells. These data imply that CTs actively interact with other surrounding cells and regulate their behavior during regeneration. Indeed, scRNA-seq revealed that during limb regeneration, PRRX1+ CT cells exhibited minor interaction in the uninjured limb tissue with other cells. Upon amputation, predicted interactions between CTs and other cell types, including MuSCs are massively increased. These newly added interactions may regulate the adaptation of surrounding cells to the reduction of CT cells.

Overall, our study established a tBid mediated conditional cell ablation system and tool in axolotls, and explored the role of CTs.

## Materials and Methods

### Axolotl care and operation

The *d/d* axolotl strain of *Ambystoma mexicanum* used in this study was originally originated from Ambystoma Genetic Stock Center (https://ambystoma.uky.edu/genetic-stock-center/). The animals were housed and bred in dechlorinated tap water at 18°C–20°C under a 12-hour light/dark cycle and fed daily. Before experimental procedures, including surgery, live imaging, electroporation, transplantation, EdU injection and harvest, the axolotls were anesthetized in 0.01% benzocaine solution (Sigma-Aldrich, E1501). Limb amputation was carried out at mid-upper forelimb of anesthetized axolotl and protruding bones were trimmed to allow proper blastema formation. Samples of uninjured or regenerated limbs were collected at indicated stages. Animal care in this study was conducted in accordance with Chinese animal welfare legislation, and approved by the Biomedical Research Ethics Committee of Guangdong Provincial People’s Hospital (license number: KY2023-1021-01).

### Tissue collection and cryosection

We collected limb bud or regenerated limb from fully anesthetized axolotls from transgenic and *d/d* axolotls for in situ hybridization and immunofluorescence detection. Isolated tissues were fixed with freshly prepared MEMFA for 1 hour at 4℃, and then dehydrated with 30% sucrose prepared in 0.8×PBS for 24 hours at 4℃. Finally, tissues were embedded in Tissue-Tek OCT (Sakura, 4583) with dry ice. The OCT blocks were stored in -80°C freezer until cryosection. For cryosection collection, the freezing microtome (Thermo Scientific, Cryostar NX70, USA) was set to -20℃. Cryosections of 10 µm thickness were collected and stored at -20℃ for further analysis.

### Axolotl strains

The transgenic axolotls *Caggs^EGFP/tBid^* (tgTol2(*CAGGs:loxP-EGFP-loxP-tBid*)^Fei^) were generated using a transposon-based method^[88, 89]^. The mix of transgene expression plasmid (*Tol2-CAGGs:loxp-EGFP-loxp-tBid*) and *Tol2* transposase mRNA was prepared on ice as described previously^[89]^. For a 5-nl injection volume per egg, the optimal amounts were 0.05 ng plasmid DNA and 0.25 ng *Tol2* mRNA. Freshly laid, single-cell stage axolotl eggs (genotypes: *d/d, Pax7^Cre-ERT^, and Prrx1^Cre-ERT^*) were collected for injection. The plasmid and *Tol2* mRNA complexes were loaded into a microinjection glass capillary and injected into the eggs using a Pneumatic PicoPump (WPI, PV830, USA) under a stereomicroscope. Approximately 200 eggs were injected. Surviving embryos were then allowed to develop into juveniles before screening. Knock-in efficiency was typically evaluated by assessing fluorescent reporter expression; for lines with weak fluorescence, genotyping was performed to accurately identify F0 transgenic animals. Moreover, *d/d*, two previously published lines, *Pax7^Cre-ERT^* (tm(*Pax7^t/+^:Pax7-P2A-memCherry-T2A-ER^T^*^2^*-Cre-ER^T^*^2^)^Etnka^)^[23, 89]^ and *Prrx1^Cre-ERT^* (tgSceI(*Mmu.Prrx1:TFPnls-T2A-Cre-ER^T^*^2^)^Etnka^)^[24]^ were used in this study.

### Plasmid construction

Plasmids were constructed using restriction enzyme-based cloning or Gibson assembly. Verification was conducted by Sanger sequencing (Tsingke, China). A comprehensive list of primers used for plasmid construction can be found in Table S2 Supplementary Information. For all downstream applications, plasmid purification was carried out using the EndoFree Plasmid Kit (Qiagen, 12162, Germany).

#### pCAGGs:Cherry-T2A-tBid

This plasmid was constructed via Gibson assembly using the following three fragments: the linearized vector backbone, and PCR-amplified *tBid* and *Cherry-T2A*. The *tBid* fragment was amplified from axolotl regeneration tail cDNA by PCR using primers tBid-F1 and tBid-R1 (Table S2, Supplementary Information). The *Cherry-T2A* fragment was amplified from the plasmid *pGEMT-Pax7-CDS-GAP43-Cherry-T2A-sERCreER* by PCR using primers Cherry-T2A-F and Cherry-T2A-R (Table S2, Supplementary Information). The vector backbone was prepared from the circular *pCAGGs:GFP-T2A-Cherry* plasmid linearized with restriction enzymes BamHI and NheI. The *Cherry-T2A* and *tBid* fragments were subsequently inserted into the linearized vector according to the manufacturer’s instructions of Gibson Assembly Kit (NEB, E2611, USA).

#### Tol2-CAGGs:loxp-EGFP-loxp-tBid

This construct was used for creation of the *Caggs^EGFP/tBid^* transgenic line. The vector backbone was prepared from the plasmid *Tol2-CAGGs:loxp-EGFP-loxp-mCherry* linearized with HindIII and NruI. The *tBid* fragment was amplified from axolotl regenerating tail cDNA by PCR using primers tBid-F2 and tBid-R2 (Table S2, Supplementary Information), and subsequently inserted into the linearized vector backbone according to the manufacturer’s instructions for the Gibson Assembly Kit.

### Tail muscle tissue electroporation

Electroporation was conducted as previously described^[90, 91]^. Axolotls were first anesthetized in 0.03% benzocaine. Subsequently, the mix of 2 μg/μl *pCAGGs:Cherry-T2A-tBid* and 2 μg/μl *pCAGGs:GFP* plasmids were mixed in 0.8×PBS containing 0.05% Fast Green dye for visual tracking, whereas the control group received a mixture of *pCAGGs:Cherry* and *pCAGGs:GFP* plasmids. The plasmid solution was microinjected into the lateral tail region posterior to the cloaca using calibrated glass micropipettes. Electroporation was performed immediately after injection in ice-cold PBS using an electroporator (Nepagene, NEPA21, Japan) equipped with 10-mm paddle electrodes (Nepagene, CUY650P10, Japan), with the following parameters: a 70 V poring pulse (5 ms) followed by four 40 V bipolar transfer pulses (50 ms) at 999 ms intervals, with a 10% voltage decay. After the procedure, animals were returned to dechlorinated tap water for recovery.

### Transplantation of blastema

We selected *Pax7:ER^T2^CreER^T^*^2^*/CAGGs:EGFP-STOP-tBid* double transgenic line as donors and *d/d* as hosts (3-4 cm in body length) in blastema transplantation experiments. Before transplantation, the upper limbs of donor animals were amputated and allowed to regenerate a blastema for 6-8 days. The donor blastemas were then collected and transplanted to the freshly amputated upper limb stumps of size-matched recipients. One months after limb blastema transplantation (regenerating complete limb), the success of limb blastema transplantation was defined by donor-derived GFP expression in the regenerated limb.

### Tamoxifen (4-OHT) and EdU administration

4-OHT (Sigma-Aldrich, H-7904) was prepared as a 20 mM stock solution in DMSO. For PAX7+ cells ablation in chimeric axolotls post-blastema transplantation, axolotls were administered intraperitoneal injections of 4-OHT at a dose of 30 mg/g body weight, given twice with a 3-day interval between injections. Juvenile axolotls (stage 38) were subjected to PAX7+ cell ablation via three 24-hour interval immersions in 2 µM 4-OHT. For CT cells ablation during development, hatched larvae were exposed to 4-OHT under two distinct treatment regimens: either 2 µM 4-OHT three times during early development (stages 36-38) or 3 µM 4-OHT nine times during later stages (stages 51-52), with administrations separated by 24-hour intervals. To investigate the effects of CT cell ablation on axolotl limb regeneration, axolotls at stage 51-52 were treated with 3 µM 4-OHT nine times, each separated by one day. Limb amputation was then performed on the tenth day following the initiation of treatment to assess regenerative outcomes. For EdU incorporation, anesthetized axolotls were treated with a single intraperitoneal injection of EdU (Thermo Scientific, A10044), at a dose of 10 μg/g of body weight. The animals were incubated for 3 hours before sample collection. All collected samples were fixed in 1 × MEMFA.

### Immunohistochemistry and EdU staining

Immunohistochemistry on cryosections was performed as previously described.^[23]^ For samples requiring EdU detection, EdU staining was performed first using the Click-iT EdU kit (Invitrogen™, C10340) according to the manufacturer’s instructions. In brief, after washing in PBS and permeabilization with PBST, slides were blocked in 5% serum prepared in PBST, and then incubated with the following primary antibodies: Chicken-anti-EGFP (Abcam, ab13970), Rat-anti-CHERRY (Invitrogen™, M11217), mouse-anti-PAX7 (DSHB), rabbit-anti-PRRX1 (a kind gift from Prof. Elly M. Tanaka’s laboratory), and rabbit-anti-Cleaved Caspase-3 (CST, 9661s). After sequential washes with PBST, nuclei were stained with DAPI (Sigma-Aldrich, D9542), and secondary antibodies included Alexa Fluor 647-donkey anti-mouse (Jackson, 715-606-150), Alexa Fluor cy3-Goat anti-Rabbit (Jackson, 711-547-003), Alexa Fluor 647-donkey anti-rabbit (Invitrogen™, A31573), DyLight550-donkey anti-rat (Invitrogen™, SA5-10027) Alexa Fluor 488-donkey anti-chicken (Jackson, 703-545-155). Finally, sections were mounted with Mowiol 4-88 (Sigma-Aldrich, 9002-89-5) mounting medium after several washes in PBST.

### Alcian blue and Alizarin red staining

Fully anesthetized axolotls were fixed in freshly prepared 4% paraformaldehyde (PFA) for 24 hours at 4℃. Following fixation, samples were washed in 0.8×PBS, and the internal organs were carefully removed. The samples were then dehydrated through a graded ethanol series (25%, 50%, and 70%). For cartilage visualization, tissues were stained with 0.01% Alcian blue 8GX (in 3:2 ethanol:glacial acetic acid) for 48 hours. After staining, samples were washed in 80% ethanol and subsequently rehydrated through a reverse ethanol series (70%, 50%, 25% and distilled water).Enzymatic digestion was then performed using 1% trypsin (MP Biomedicals, #153571) at 37℃ for 1 hour. Post-digestion, samples were rinsed in 1% KOH and stained with 0.01% Alizarin Red (in 1% KOH) for bone visualization. Samples were then cleared by repeated washing with 1% KOH until transparent. Finally, the samples were dehydrated through a graded ethanol series (25%, 50%, 70%, 90% and 100%), followed by transition through graded glycerol/ethanol solutions (1:3, 1:1 and 3:1), and stored long-term in 100% glycerol at room temperature. Imaging was performed using an Olympus SZX10 microscope (Olympus, Tokyo, Japan).

### In situ hybridization

Total RNA was extracted from regenerating axolotl limb tissues at various stages and reverse-transcribed into cDNA. Target genes were amplified from this cDNA using oligonucleotide primers containing a T7 promoter sequence at the 5′ end (Table S3, Supplementary Information). The PCR products were used as templates for in vitro transcription to generate DIG-labeled antisense RNA probes.

Tissue sections were washed with PBST and incubated in hybridization buffer containing 550 ng/mL of DIG-labeled *Hoxa9*, *Hoxa11*, or *Hoxa13* RNA probes overnight. Following hybridization, the sections were washed with SSC buffer and treated with RNase A (Sigma-Aldrich, R4642) to remove non-specific probes. Finally, sections were incubated with anti-Digoxigenin-AP antibody (Roche, 11093274910) and developed with BM Purple substrate (Roche, 11442074001) to visualize colored precipitates at specific tissue sites. The reaction was stopped with 1 mM EDTA in PBS.

### Imaging

Bright field and fluorescence imaging of live axolotls were captured using an Olympus stereomicroscope (SXZ16). Immunohistochemistry and in situ hybridization signals in cryosections were acquired using a laser-scanning confocal microscope (Zeiss, LSM980) and an inverted microscope (Olympus, IX83), respectively.

### Single cell dissociation of scRNA-seq

Regenerating limb tissues were collected from *Caggs^EGFP/tBid^* control and *Prrx1-tBid dTG* axolotls following 4-OHT treatment, spanning from approximately 500 µm proximal to the amputation plane at three time points: 6, 10, and 15 dpa. Considering that the experimental group underwent amputation at upperarm, forelimb samples from control axolotls treated with 4-OHT six times were harvested exclusively from the upper limb region to serve as 0 dpa controls. After a brief rinse with 0.8× PBS, the limb samples were minced using sterile surgical scissors and transferred into 1 mL of dissociation solution containing 1× Liberase, 0.1 U·µl ^-1^ DNase I in 0.8× PBS. The tissues were digested at 37℃ in a water bath for approximately 20 minutes, with gentle shaking every 10 min. The digestion was terminated by adding 10% FBS. The resulting cell suspension was filtered through a 70 μm cell strainer to remove tissue debris, and cells were collected by centrifugation at 500 g, 4℃ for 5 minutes. The cells were resuspended in 2 mL of 0.8× PBS and filtered through a 30 μm cell strainer. Following a second centrifugation under the same conditions, the cells were resuspended in 0.5 mL of 0.8× PBS to generate a single-cell suspension for subsequent library construction.

### Construction and sequencing of single-cell RNA-seq libraries

Single-cell RNA-seq libraries were constructed using the SeekOne Digital Droplet Single Cell 3’ Library Preparation Kit (SeekGene, Cat. No. K00202) according to the manufacturer’s instructions. Briefly, mixing of cells with reverse transcription reagents was performed prior to loading into the SeekOne Chip S3 sample well. Barcoded hydrogel beads and partitioning oil were dispensed into the designated wells to generate emulsion droplets, followed by reverse transcription (42℃, 90 min) and enzyme inactivation (85℃, 5 min). Next, library preparation, including cleanup, fragmentation, end repair, A-tailing, and adapter ligation, was performed on the amplified cDNA. To amplify DNA fragments representing the 3’ poly(A) regions of expressed genes, an indexed PCR was conducted to incorporate both cell barcodes and unique molecular identifiers (UMIs). Following purification and quality control (Qubit and Bio-Fragment Analyzer), the final libraries were sequenced on an Illumina NovaSeq 6000 (PE150).

### Generation of the scRNA-seq data

The raw reads were processed with Fastp^[92]^ to trim primer sequences and perform quality filtering. The resulting high-quality reads were then used for downstream analysis. The cleaned reads were processed using the SeekSoul Tools (https://github.com/seekgenebio/seeksoultools) pipeline to generate a transcript expression matrix. Initially, cell barcodes and UMIs were extracted according to a predefined structure specifying the positions of the barcode, linker, and UMI in each read. The barcodes were then corrected against a whitelist. The corrected barcodes and corresponding UMIs were embedded into the read headers. Subsequently, the reads were aligned to the reference genome (AmexG_v6.0_DD)^[93]^ with STAR (v2.7.10a)^[94]^. After the alignment procedure mapped the reads to the genome, QualiMap was used in conjunction with a gene annotation file (GTF) to assign the reads to exonic, intronic, and intergenic regions. Additionally, featureCounts was employed to annotate the genome-aligned reads to genes. SeekSoul Tools employed the adjacency algorithm from UMI-tools to consolidate UMIs exhibiting high sequence similarity (typically differing by a single nucleotide) and substantial count disparities, whereby low-count UMIs were merged into their corresponding high-count analogues. Finally, all barcodes were sorted based on UMI counts to retain confidently identified cells, resulting in a filtered expression matrix for downstream analyses.

### The processing of scRNA data

Data quality control, preprocessing, and dimensionality reduction were performed using Seurat (v4.1.0)^[77]^. Low-quality cells (expressing <500 genes and with >5% mitochondrial gene content) were excluded. Doublet removal was performed using DoubletFinder (v2.0.3)^[78]^, discarding the top 5% of predicted doublets per library. Expression matrices were normalized using the NormalizeData function. Highly variable gene were identified based on dispersion, with the top 2,000 most variably expressed genes selected for subsequent analysis. Linear dimensionality reduction was performed using principal component analysis (PCA), and the top 30 principal components, capturing the most significant biological variation, were retained for downstream clustering analyses. To integrate expression data across samples and mitigate batch effects, a reciprocal PCA (RPCA)-based integration approach was applied. Cell clusters were subsequently identified using a graph-based clustering algorithm with a resolution parameter of 0.5. To visualize the data structure, Uniform Manifold Approximation and Projection (UMAP) was applied to reduce dimensionality, preserving the global topology of the single-cell dataset in a two-dimensional embedding.

### RNA velocity

The RNA velocity was performed by Dynamo (v1.3.2) following the instructions at https://github.com/aristoteleo/dynamo-release^[81]^. Unspliced and spliced RNA count matrices were produced using Velocyto^[95]^, based on the aligned sequencing data (BAM file) and genomic annotations. The unspliced and spliced count matrices from regeneration-related cell types were subjected to preprocessing with dyn.pp.Preprocessor for normalization and highly variable gene selection. The dyn.tl.dynamics function was used to infer mRNA transcription and degradation kinetics based on a stochastic dynamical model. Then, instantaneous RNA velocity vectors were computed for individual cells through Pearson correlation using the dyn.tl.cell_velocities function, and their confidence levels were assessed with dyn.tl.cell_wise_confidence. Finally, high-confidence velocity vectors were projected onto the UMAP embedding to construct streamline plots that visualize potential cell-state transitions.

### Gene module analysis

To investigate the migration of newborn CT cells during regeneration, differentially expressed genes from distally and proximally amputated regenerative blastema were collected and used as predefined gene sets. Gene module expression scores were computed to quantify the activity of predefined gene sets in each cell. Proximal and distal enrichment scores for each cell were calculated using the AddModuleScore function from the Seurat (v4.1.0)^[77]^ with default parameters (ctrl = 100, nbin = 24). The complete gene lists used in this analysis are provided in Table S1 Supplementary Information.

### Cell-Cell communication analysis

To investigate intercellular communication networks during axolotl limb regeneration, specifically examining interactions between CT cells and other cell types, as well as differential cellular interactions between *Prrx1-tBid dTG* and *Caggs^EGFP/tBid^* axolotls, the CellChat (v2.2.0)^[96]^ package was used. Normalized expression data and cell cluster annotations were retrieved from the seurat object using the GetAssayData function. A CellChat object was initialized using the human secreted signaling ligand-receptor interactome database (CellChatDB.human, Secreted Signaling). Enriched ligands, receptors, and ligand-receptor (LR) pairs within the dataset were identified and projected onto a curated protein-protein interaction (PPI) network. Communication probabilities were computed using the computeCommunProb function and subsequently filtered with filterCommunication. The aggregated network was then generated using the aggregateNet function to summarize communication probabilities and interaction weights across all signaling pathways. To compare the differences in cellular interactions across regeneration time points in control axolotls, specifically comparing *Prrx1-tBid dTG* with control axolotls, we first generated CellChat objects for each stage separately and subsequently merged different CellChat objects using the mergeCellChat function. Finally, visualization was performed using the netVisual_diffInteraction and netVisual_heatmap functions.

### Quantifications and statistical analyses

All quantifications were performed manually. For immunohistochemistry-based cell counts, one 10µm thick section was collected from every eight consecutive slices. Limb samples were sectioned longitudinally. To assess PRRX1+ cell ablation during development, all target cells were counted across approximately 4-5 sections per sample. For assessing PRRX1+ cell ablation in regeneration, quantification included all target cells within 5-8 sections encompassing both the newly regenerated blastema and the 500 µm region proximal to the amputation plane.

All data are presented as mean ± SEM. Statistical comparisons between experimental groups were performed using an unpaired t-test in GraphPad Prism 9.0.0. A p-value less than 0.05 was considered statistically significant. Statistical significance in figures is indicated as follows: n.s. (*P* > 0.05), * (*P* < 0.05), ** (*P* < 0.01), *** (*P* < 0.001), **** (*P* < 0.0001).

## Supporting Information

Supporting Information is available from the Wiley Online Library or from the author.

## Acknowledgments

We extend our gratitude to Guoqing Liu, Zhenyao Wu, and Yingqiao Tang for their diligent care and maintenance of the axolotls. This work was supported by the following grants: National Key R&D Program of China (2023YFA1800600 and 2021YFA0805000 to JF.F.); National Natural Science Foundation of China (32300701 to Z.H.; 92268114 to JF.F.; 32070819 to Y.L.); High-level Hospital Construction Project of Guangdong Provincial People’s Hospital (DFJHBF202103 and KJ012021012 to JF.F.); GuangDong Basic and Applied Basic Research Foundation (2024A1515110284 to W.F.)

## Author contributions

Y.H, W.F., Z.H., and J.Z. contributed equally to this work. Y.H, W.F., and Z.H. designed and performed the experiments, as well as analyzed the results. Y.H. and W.F. wrote the original draft. J.Z., J.T., S.L., T.Y., H.M., B.L. and S.F. contributed to the experimental work and data analysis. JF.F. and Y.L. conceived and designed the study, supervised the project, and revised the manuscript. All authors reviewed and approved the final manuscript.

## Data availability Statement

The data that support the findings of this study are available from the corresponding author upon reasonable request. The scRNA-seq raw data in this study are openly available in the National Genomics Data Center [https://ngdc.cncb.ac.cn/] under the BioProject number PRJCA052336.

